# Divergence of Bowman-Birk Protease Inhibitor Family into seed-specific and environmentally responsive subfamilies in Legume and Soybean: Implication for Legume Seed Composition Improvement

**DOI:** 10.1101/2025.08.26.658694

**Authors:** Zhibo Wang, Helen Jiang, Keshun Liu, Neeta Lohani, Saurav Misra, Whenhao Shen, Luis Gomez-Luciano, Suresh Pokhrel, Ray Collier, Shawn M. Kaeppler, Yong-Qiang Charles An

**Affiliations:** Donald Danforth Plant Science Center, St. Louis, MO, USA; U.S. Department of Agriculture, Agricultural Research Service (UDA-ARS), National Small Grains and Potato Research Unit, Aberdeen, Idaho, USA; USDA-ARS, Plant Genetics Research Unit, St. Louis, MO, USA; Department of Biochemistry and Molecular Biophysics, Kansas State University, Manhattan, KS, USA; Molecular Technologies Department, Wisconsin Crop Innovation Center, University of Wisconsin-Madison, Madison, WI, USA

## Abstract

Bowman-Birk inhibitors (BBI) are an ancient class of serine protease inhibitors originating prior to the emergence of the angiosperms. While BBIs have been preserved in the legume (Fabaceae) and cereal (Poaceae) families, they have been lost in many other divergent lineages. However, their underlying molecular evolution and regulation of BBI remain largely uncharacterized. Our study shows that BBIs in legumes and cereals are encoded by two large and divergent gene families. BBI genes in legumes have further diversified into two subfamilies with distinct gene expression patterns. Genes in one legume BBI subfamily are specifically expressed in seeds while BBI genes in the other legume subfamily and cereal do not have significant expressions in any examined tissues including seed, root, leaf and flower. The soybean BBI gene family shows evidence of expansion via whole genome, segmental and tandem duplication. Protein sequence and structural analysis predicts that functional domains for double-headed inhibitory loops and binding abilities to trypsin and chymotrypsin are largely preserved within the soybean BBI family. The seed-specific subfamily genes are specifically expressed at maturation stages and not at embryogenesis stages. The other, non-seed BBI subfamily genes are highly responsive to a distinct spectrum of signals related to abiotic and biotic stresses. Their specific expression under non-essential biological processes for plant growth and development suggests that, although BBIs have been retained in both cereals and legumes, likely due to their role in enhancing plant fitness under natural selection pressures, they are not involved in core developmental processes. This may explain why BBIs were lost in many divergent plant lineages and support their well-established roles in plant adaptation to environmental stress. Having knocked out the seed-specific BBIs through a CRISPR/Cas9 approach, we have successfully generated soybeans which exhibited 69.4 - 73.7% reduction of trypsin inhibitor activity and 76.4 - 79.4% reduced chymotrypsin inhibitor activity. The edited soybean did not show significant changes in key agronomic traits, supporting that the functions of BBIs are not essential. While BBIs in soybean seeds may have a desirable function in natural selection, they are antinutrients from an applied perspective for their use in feed and food. It provides an opportunity to reduce BBIs in seeds for quality improvement. Our findings provide insights into molecular evolution, regulation, and function of BBI in plants, and successfully demonstrate engineering BBI in seeds to result in production of food and feed of higher nutritional value with minimal impacts on the agronomic performance of the plant.

## Introduction

Bowman-Birk inhibitors (BBI) are an extensively studied family of serine protease inhibitors in plants. The prototypical BBI was first discovered in soybeans (*Glycine max*) 80 years ago (Bowman, 1946). Subsequently, BBIs have been identified in divergent plant species, most prominently in legumes (*Fabaceae*), including soybean, peanut, and burclover, and in cereals (*Poaceae*), including rice, maize, and sorghum (Mello *et al*., 2003; Qi *et al*., 2005). The genome of *Selaginella moellendorffii,* a member of the lycopod plant lineage that diverged 200 to 230 million years ago, was recently reported to contain BBI genes, suggesting that the BBI family is ancient and arose prior to the emergence of angiosperms (James *et al*., 2017). However, no BBI gene was identified in several angiosperm lineages, including Spirodela *polyrhiza* in monocots and *Malpighiales* and *Asterids* in eudicots (James *et al*., 2017). In the Rosid clade of eudicots, Arabidopsis and cucumber also lack BBI genes (James *et al*., 2017). Evolutionary pressures that contribute to the preservation of BBI gene families in agriculturally important legumes and cereals are poorly understood.

BBIs and their biochemical characteristics have been extensively studied in legume species (De Paola *et al*., 2012; Lioi *et al*., 2009; Ragg *et al*., 2006). Legume BBIs are dual inhibitors of both trypsin and chymotrypsin (Birk, 1961; Birk *et al*., 1963). This duality is attributed to two separate inhibitory loops of BBIs, each stabilized by a disulfide bridge (Odani and Ikenaka, 1972). These two spatially separated loops give rise to a “double-headed” structure, predicted to have arisen from an internal gene duplication of a single-headed ancestral inhibitor (Mello *et al*., 2003). Although BBIs in both cereals and legumes share a common and ancient ancestor (Mello *et al*., 2003), cereal BBIs lack the double inhibitory-loop structure common in legume BBIs, due to the absence of inhibitory-loop-forming cysteine residues, rendering the second loop nonfunctional (Park *et al*., 2004; Qi *et al*., 2005).

Increasing evidence indicates that expression and functions of BBIs are subjected to a complex genetic regulation. BBIs are often encoded by gene families with various sizes across different plant species (James *et al*., 2017; Mello *et al*., 2003). For instance, *Phalaenopsis equestris* has only one BBI gene, while another monocot species, *Musa acuminata*, has ten BBI genes (James *et al*., 2017). They are also associated with diverse biological roles, including antimicrobial activity. For instance, the rice BBI OsBBI5 enhances OsAPIP5-mediated programmed cell death (PCD), conferring broad-spectrum resistance to *Magnaporthe oryzae* and *Xanthomonas oryzae* pv. *Oryzae* (Zhang *et al*., 2024). In wheat, genetic mapping identified putative BBI genes as candidates for seedling resistance to tan spot and Fusarium head blight (Juliana *et al*., 2018; Sari *et al*., 2019). BBIs also contribute to abiotic stress tolerance, including salinity, oxidative stress, and drought, though the underlying mechanisms, such as their role in iron uptake regulation, remain unclear. In wheat, the BBI-type gene WRSI5 enhances salt tolerance (Shan *et al*., 2008), while in peanuts, a BBI gene is upregulated by water deficit and jasmonic acid (JA) but repressed by abscisic acid (ABA) after 24 hours of treatment (Drame *et al*., 2013). Wheat BBI gene expression also changes in response to heat and drought stress (Xie *et al*., 2021). Due to their ability to inhibit protease activity and reduce protein digestion efficiency, BBIs can deter herbivores from consuming seeds (Sultana *et al*., 2023; Wang *et al*., 2015). High accumulation of BBI is often detected in seeds, and BBIs have been considered to be sulfur storage proteins and a rich source of sulfur-containing amino acids due to high cysteine content (Birk, 1985). However, BBI is a major antinutrient that significantly reduces the nutritional value of soybeans, posing a significant challenge in the current soybean industry. To date, the molecular regulation of BBI production and the role of BBIs in plant growth and development, particularly in seed development, remain poorly understood. Understanding their underlying molecular regulation and biological functions of BBIs in plant development is crucial for the development of effective strategies to reduce BBI levels in seeds and the improvement of soybean nutritional value without negative impact on soybean agronomic performance.

In this study, we comprehensively investigated genes encoding BBIs in legumes and cereals and their evolution. We found that legume and cereal BBI genes have diverged into two distinct families, with legume BBI genes further diverged into a seed-specific and a highly environmental signal responsive subfamily. We found that the soybean BBI gene family expanded through diverse mechanisms including whole genome, segmental and tandem duplication. However, structural analysis and protein modeling shows that soybean BBIs have largely preserved their functional domains, including ER signal sequences, target protease binding determinants and binding activity. These findings suggest that most BBI genes encode functional BBI proteins that reflect the functional constraints and characteristics of the prototypical family members. We applied a CRISPR/Cas9 system to selectively knock out seed-specific BBI genes in soybeans while preserving the highly responsive BBI genes. The edited soybeans have significantly reduced protease inhibitor activities, but do not exhibit significant changes in agronomic traits. Therefore, our study provides a foundation for breeding low-BBI soybean and legume cultivars with high nutritional value without significantly compromised agronomic performance.

## Methods and Materials

### Plant growth and greenhouse conditions

*Glycine max* cv. Williams 82 (WM82) and transgenic plants derived from the WM82 background were cultivated in a Donald Danforth Plant Science Center greenhouse, each in one-gallon pots filled with Berger BM7 bark soil (10121500; Hummert, Earth City, MO, USA). The plants were watered twice daily by an automatic irrigation system and fertilized using a 15-16-17 nitrogen-phosphate-potash formula. Supplemental light was provided to ensure a 14h/10h light/dark cycle at 25°C/20°C with a relative humidity of >□40%.

### Identification and phylogeny of BBI genes

Genes containing the BBI domain (Pfam accession PF00228) were identified in 11 species: six *Fabaceae*, four *Poaceae*, and Cinnamomum kanehirae (CKAN) as an outgroup species (Table S1). These genes sequences were obtained from the Phytozome database (https://phytozome-next.jgi.doe.gov), except for *Phaseolus vulgaris*, *Vigna unguiculata*, *Medicago truncatula*, and *Arachis hypogea* which were obtained from the Legume Information System database (Berendzen *et al*., 2021). Coding sequences were aligned in MAFFT (Katoh and Standley, 2013) using the L-INS-i method with 1000 maximum iterations, and a maximum likelihood tree was generated using the GTR+F+I+G4 model in IQ-TREE (Nguyen *et al*., 2015) with a 1000 ultrafast bootstrap approximation.

### Tissue-specific expression analyses of BBIs in two cereal and four legume plants

To determine expression patterns of BBI genes in *Glycine max*, *Glycine soja*, *Vigna unguiculata*, *Oryza sativa* and *Zea mays*, RNA-Seq data sets were downloaded from the NCBI Sequence Read Archive database (Table S2). The gene expression data for *Phaseolus vulgaris* was downloaded from the Common Bean Gene Expression Atlas (https://www.zhaolab.org/PvGEA/page/download). The ComplexHeatmap package was used to visualize expressions of the genes of interest on a heat map.

### Analysis of soybean BBI transcriptional response to different environmental treatments

We applied a transcriptome platform developed in the laboratory (Shen, et.al, unpublished) to determine transcriptional expression of BBI genes in response to different internal and external signals. Briefly, a consolidated collection of RNA-Seq datasets were used in the study. Differentially expressed genes (DEG) were identified by comparing transcriptional accumulation of each BBI gene for each treatment to its corresponding control. The treatments for each signal, such as all drought treatments or combinations of multiple signals such as drought + mepiquat chloride, were grouped together to represent soybean response to the signal. The total number of treatments in a signal group that differentially regulated a BBI gene were calculated and visualized in a heatmap.

### Synteny and Gene duplication Analysis

Synteny detection and visualization was performed using MCScanX (Tang *et al*., 2008). Gene duplications across the BBI genes in *G. max* were identified by performing an all-by-all protein gene duplication analysis using the DupGen_finder pipeline (Qiao *et al*., 2019). This pipeline identified different modes of gene duplication in plants: whole-genome, tandem, proximal, transposed, or dispersed duplication. For visualization of the gene duplications, the chromosomal locations of the genes were plotted using the IGV (Integrated Genomics Viewer) (Thorvaldsdottir *et al*., 2013) and adapting a color scheme to represent gene duplications.

### Multiple Sequence Alignment and Protein Structure

MUSCLE (Edgar, 2004) was used to align BBI protein sequences to highlight their BBI domains. We further generated structural models of the GmBBI (*Glycine max* BBI) sequences using a Google colab version of AlphaFold2_mmseqs2 (ColabFold v. 1.5.5) current as of July 1, 2024 (Mirdita *et al*., 2022). We adjusted the AlphaFold2 settings to provide the top 5 amber-relaxed models, with the following options set at values other than their defaults: template_mode: pdb100; use_dropout: checked. The sequences of bovine trypsin and chymotrypsin were used to generate the respective BBI:protease complexes (Uniprot accession codes: P00760 and P00766). We compared the generated structures to selected experimental NMR and X-ray crystallography structures of apo-BBI1 from the RCSB including PDB accessions 1k9b (Voss *et al*., 1996), 1bbi and 2bbi (Werner and Wemmer, 1992). We compared the generated BBI:protease complex models to X-ray structures of BBI1 complexed with trypsin and chymotrypsin, PDB accessions 1d6r (Koepke *et al*., 2000) and 5j4q (Tornøe *et al*., 2017). All structural visualizations, comparisons and figure generation were carried out using open-source PyMol (The PyMOL Molecular Graphics System, Version 3.0, Schrödinger LLC).

### CRISPR/Cas9 vector construction and soybean transformation

A CRISPR/Cas9 vector, pSp-Cas9-BBI, was designed and assembled in the WCIC-MTD DICOT RK2 synthetic binary plasmid backbone at the Molecular Technologies Department within the Wisconsin Crop Innovation Center at the University of Wisconsin – Madison. Four guide RNAs (gRNAs) targeting eight *BBI* genes were designed using CRISPRdirect software (https://academic.oup.com/bioinformatics/article/31/7/1120/180346). gRNAs containing off-targets involving one or more nucleotides mismatches were filtered out based on Soybean (*Glycine max*) genome, v2.0. and only BBI genes-specific gRNAs were used. A polycistronic gRNA expression cassette, using the tRNA_Gly_ approach (Xie *et al*., 2015), was assembled as a MoClo (Weber *et al*., 2011) compatible Level 0 CDS1 Golden Gate part using the WCIC-MTD BLACKSMITHv4 system. The gRNA cassette was used, in a Level 1 Golden Gate reaction, to produce a transcriptional unit (TU) under control of the Cestrum Yellow Leaf Curling virus promoter (*CmYLCV*p, (Stavolone *et al*., 2003)), with polyadenylation provided by the *Nicotiana benthamiana Actin 3* terminator (*NbACT3*t, (Diamos and Mason, 2018)). Importantly, since the gRNA cassette was under control of a DNA dependent RNA Pol II promoter and terminator, which does result in production of a polyadenylated mRNA, a tRNA_Gly_ was included in the polycistronic gRNA expression cassette downstream of the final gRNA scaffold, and upstream of the polyadenylation signal, to ensure that the poly A tail would be cleaved away from gRNA #4 to ensure that this gRNA could bind with Cas9. The *Glycine max Ubiquitin 3 XL* promoter (*GmubiXL*p, (De La Torre and Finer, 2015)) was used to drive expression of the *SpCas9* which had been codon optimized for soybean (*GmCas9*, (Michno *et al*., 2015)), and which was additionally intronized with the *Solanum tuberosum LS1* intron (Soltu.DM.07G028260, (Eckes *et al*., 1986; Vancanneyt *et al*., 1990)); polyadenylation was under control of the *Glycine max Ubiquitin 1* terminator (*GmUbi1*t (Glyma.10g251900.1), (Xia *et al*., 1994)). The plant selectable marker transcriptional unit, which confers to transgenic plant cells the ability to survive spectinomycin selection, was comprised of a 2X enhanced ((Kay *et al*., 1987)) Cauliflower Mosaic Virus 35S promoter (2XCaMV35Sp, (Odell *et al*., 1985)), additionally outfitted with the translational enhancer from Tobacco Mosaic Virus (Gallie, 2002; Gallie and Walbot, 1992)), to control expression of a fusion protein comprised of the dual organellar (mitochondria and plastids) targeting peptide from the *Arabidopsis thaliana Threonyl tRNA Synthetase* (AtdTP from At2g04842; (Berglund *et al*., 2009)) fused in frame with the *Escherichia coli aminoglycoside 3”-adenyltransferase* (*aadA1a*, (Hollingshead and Vapnek, 1985)), with polyadenylation under control of the Cauliflower Mosaic Virus 35S terminator (Hirt *et al*., 1990; Irniger *et al*., 1992). Finally, to facilitate non-destructive identification of null segregants in the T_1_ progeny to support finding edited individuals from which the transgenes had been eliminated via segregation, the construct also included the SeedRUBYv1 marker (similar to the previously reported RUBY marker (He *et al*., 2020), but assembled using the WCIC-MTD RIGGERv1 system for Golden Gate mediated assembly of polycistronic messages under control of the *GmScream6* seed specific promoter (Glyma.10g39150, (Zhang *et al*., 2015)), with polyadenylation under control of the intron-less *Extensin* terminator from *Nicotiana tabacum* (LOC107768502, (Rosenthal *et al*., 2018)). All Level 1 transcriptional units (TUs) were assembled into a Level 2 Golden Gate reaction (see Supplemental Methods) using the WCIC DICOT RK2 binary plasmid backbone, yielding the final construct: plasmid stock number RC6458A-RCGG6180 (also referred to as BBI SOYEDIT6). The construct was transformed via electroporation (McCormac *et al*., 1998) into *Agrobacterium rhizogenes* strain 18r12v (produced in 2004 by Collier and Taylor, unpublished), which is a disarmed version of *Agrobacterium rhizogenes* strain NCPPB2659 (Combard *et al*., 1987). The resulting Ar18r12v clone (stock number WCIC-A-01058) was used for production of transgenic WM82 via Agrobacterium-mediated transformation at the Wisconsin Crop Innovation Center. The lines from five independent transgenic events were generated and were genotyped.

### Genotyping of gene editing events

Genomic DNA was extracted from both leaf and seed tissues of plants using the CTAB method. Subsequently, each target BBI gene was amplified via PCR and sequenced for genotyping using Sanger sequencing. Details of the PCR and sequencing primers are provided in Table S3.

### Trypsin and chymotrypsin inhibition measurements and BBI protein identification by SDS-PAGE

WM82 (wild type) and edited line dry seeds were characterized as duplicates in trypsin inhibitor activity (TIA) and chymotrypsin inhibitor activity (CIA) assays. TIA in soybean seeds was measured by American Oil Chemists Society Official Method Ba 12a-2020 (Liu *et al*., 2021), and calculated as mg trypsin inhibited (TId) per g sample (Liu, 2021). CIA was determined according to the method of Liu (Liu, 2022) and calculated as mg chymotrypsin inhibited (CId) per g sample (Liu, 2023). Non-reducing Tricine SDS-PAGE was performed for seeds of Williams 82 (control) and edited lines, and the same extracts were used to measure TIA and CIA. The SDS-PAGE results were used to quantify levels of BBI proteins in seeds based on band intensity against purified BBI protein, as described previously (Liu, 2024).

### Measurements of seed composition traits

A near-infrared reflectance (NIR) analyzer (Model, DA7250 NIR Analyzer, PerkinElmer, Springfield, IL) was used to measure seed composition traits in triplicate for T2 seeds. For each sample, about 60 mL of seeds were used, and results for each quality parameter were averaged.

## Results

### The Legume BBI family has evolved to become highly divergent from cereal BBIs

We identified BBI genes containing the BBI domain (PF00228) in six legume species (*Glycine max*, *Glycine soja*, *Vigna unguiculata*, *Phaseolus vulgaris*, *Medicago truncatula*, and *Arachis hypogaea*), four cereal species (*Sorghum bicolor*, *Triticum aestivum*, *Oryza sativa*, and *Zea mays*), and an outgroup species *Cinnamomum kanehirae* (CKAN) (Table S1). The number of BBI genes was highly variable among the species (Table 1). Within the six legume species, the number of BBI genes ranged from three in *Phaseolus vulgaris* to 13 in *Glycine max*. Within the four cereal species, the number of BBI genes ranged from four in *Sorghum bicolor* to ten in *Oryza sativa*. Phylogenetic analyses show that legume BBIs and cereal BBIs respectively cluster into two separate clades (Figure 1). This clustering pattern suggests that the BBIs in each species arose from gene duplication events following divergence from the most recent common ancestor shared by the *Fabaceae* and *Poaceae*. Legume BBIs further form two distinct clades. Interestingly, one clade contains only genes from the plant tribe *Phaseoleae*, which includes *Glycine max*, *Glycine soja*, *Phaseolus vulgaris*, and *Vigna unguiculata* in addition to *Medicago truncatula* and *Arachis hypogaea* (Figure 1). The *Phaseoleae* tribe includes species currently cultivated for seeds used for both animal feed and human foods. We speculate that BBIs may contribute to characteristic traits specific to this tribe. The other legume clade contains genes from all analyzed species of legume, except for the common bean (Figure 1), which may have lost a previously present gene from this clade.

**Figure 1.**
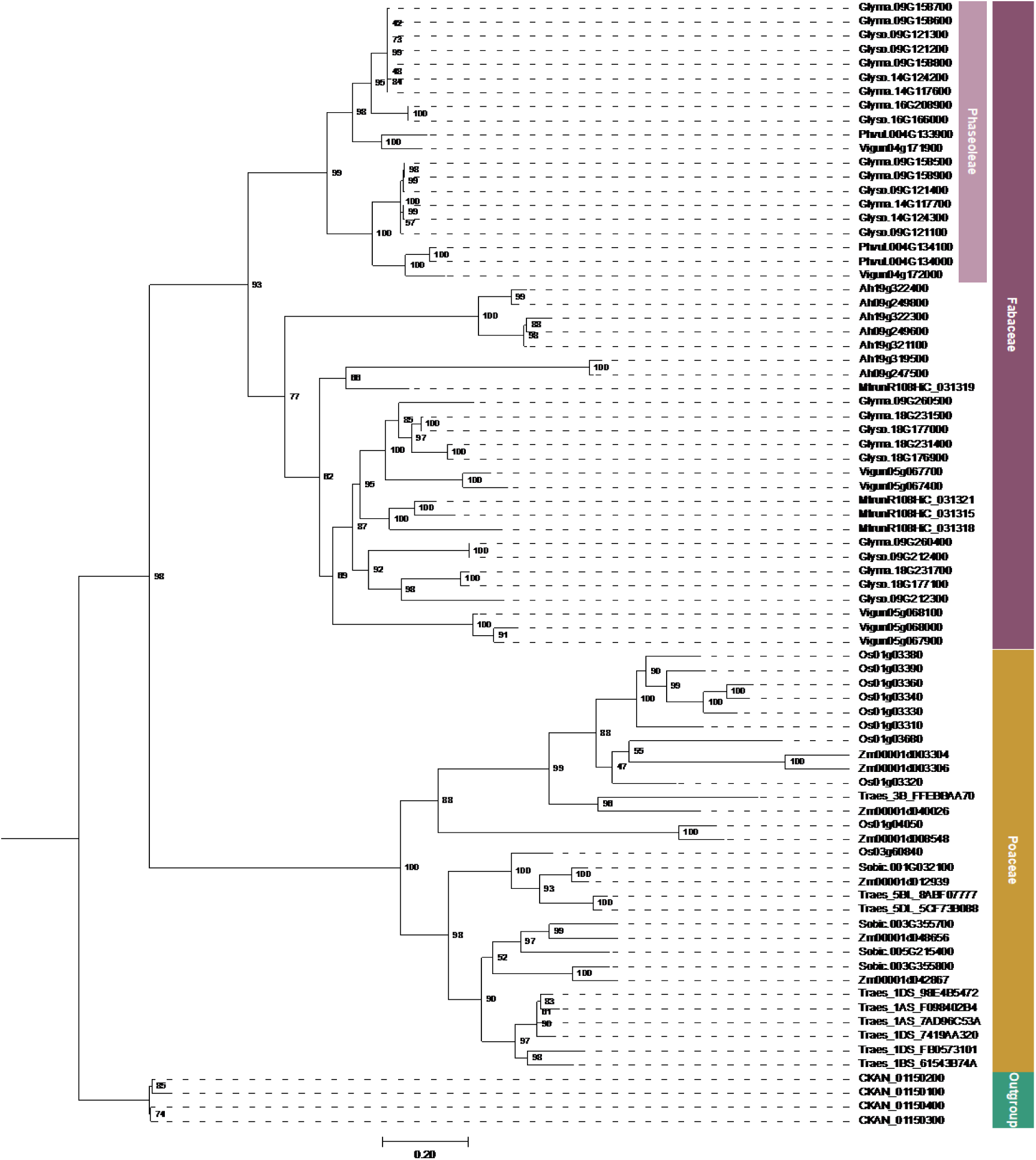
Phylogeny of BBI genes in 11 plant species. It includes four *Poaceae* species, *Sorghum bicolor* (*Sobic*), *Zea mays* (*Zm*), *Oryza sativa* (*Os*), and *Triticum aestivum* (*Traes*); six *Fabaceae* species, *Medicago truncatula* (*Mtrun*), *Phaseolus vulgaris* (*Phvul*), *Glycine max* (*Glyma*), *Glycine soja* (*Glyso*), *Arachis hypogaea* (*Ah*), and *Vigna unguiculata* (*Vigun*); and an outgroup species CKAN. The phylogenetic tree was constructed based on the evolutionary relationship.

**Table 1.** Number of BBI genes in the examined species.

|  | Common name | Latin name | No. of genes | Family | Gene abb. initials |
| --- | --- | --- | --- | --- | --- |
| 1 | Sorghum | *Sorghum bicolor* | 4 | *Poaceae* | *Sobic* |
| 2 | Corn | *Zea mays* | 7 | *Poaceae* | *Zm* |
| 3 | Rice | *Oryza sativa* | 10 | *Poaceae* | *Os* |
| 4 | Wheat | *Triticum aestivum* | 9 | *Poaceae* | *Traes* |
| 5 | Barrelclover | *Medicago truncatula* | 4 | *Fabaceae* | *Mtrun* |
| 6 | Common bean | *Phaseolus vulgaris* | 3 | *Fabaceae* | *Phvul* |
| 7 | Soybeans | *Glycine max* | 13 | *Fabaceae* | *Glyma* |
| 8 | Wild soybeans | *Glycine soja* | 12 | *Fabaceae* | *Glyso* |
| 9 | Peanut | *Arachis hypogaea* | 7 | *Fabaceae* | *Ah* |
| 10 | Cowpea | *Vigna unguiculata* | 7 | *Fabaceae* | *Vigun* |
| 11 | Stout camphor tree | *Cinnamomum kanehirae* | 4 | *Lauraceae* | *LOC* |

### BBIs evolved into two divergent subfamilies in legume species with distinct and conserved expression patterns

To gain insights into the functions of BBIs in legumes and cereals, we analyzed BBI gene expression in two vegetative organs (leaf and root) and two reproductive organs (flower and seed) from four legume species (*Glycine max*, *Glycine soja*, *Vigna unguiculata*, and *Phaseolus vulgaris*) and two cereal species (*Oryza sativa* and *Zea mays*). All BBI genes in legume clade 1 were specifically expressed in seeds (Figure 2). In contrast, BBI genes in legume clade 2 exhibited undetectable or very low expression across all examined tissues. The division of BBI genes into two clades and conservation within each clade, both in sequences and expression patterns, implies that the legume BBI subfamilies evolved to assume distinct biological functions. Interestingly, like legume BBI genes in subclade 2, all BBI genes in the cereal species, as represented by rice and maize, displayed no or very low expression in root, leaf, flower or seeds (Figure 2). Thus, conserved seed-specific expression patterns of BBI genes of the legume subclade 1, together with their specific presence in *Phaseoleae*, suggest that the seed-specific BBI genes arose more recently and evolved an important functional role in *Phaseoleae* seeds.

**Figure 2.**
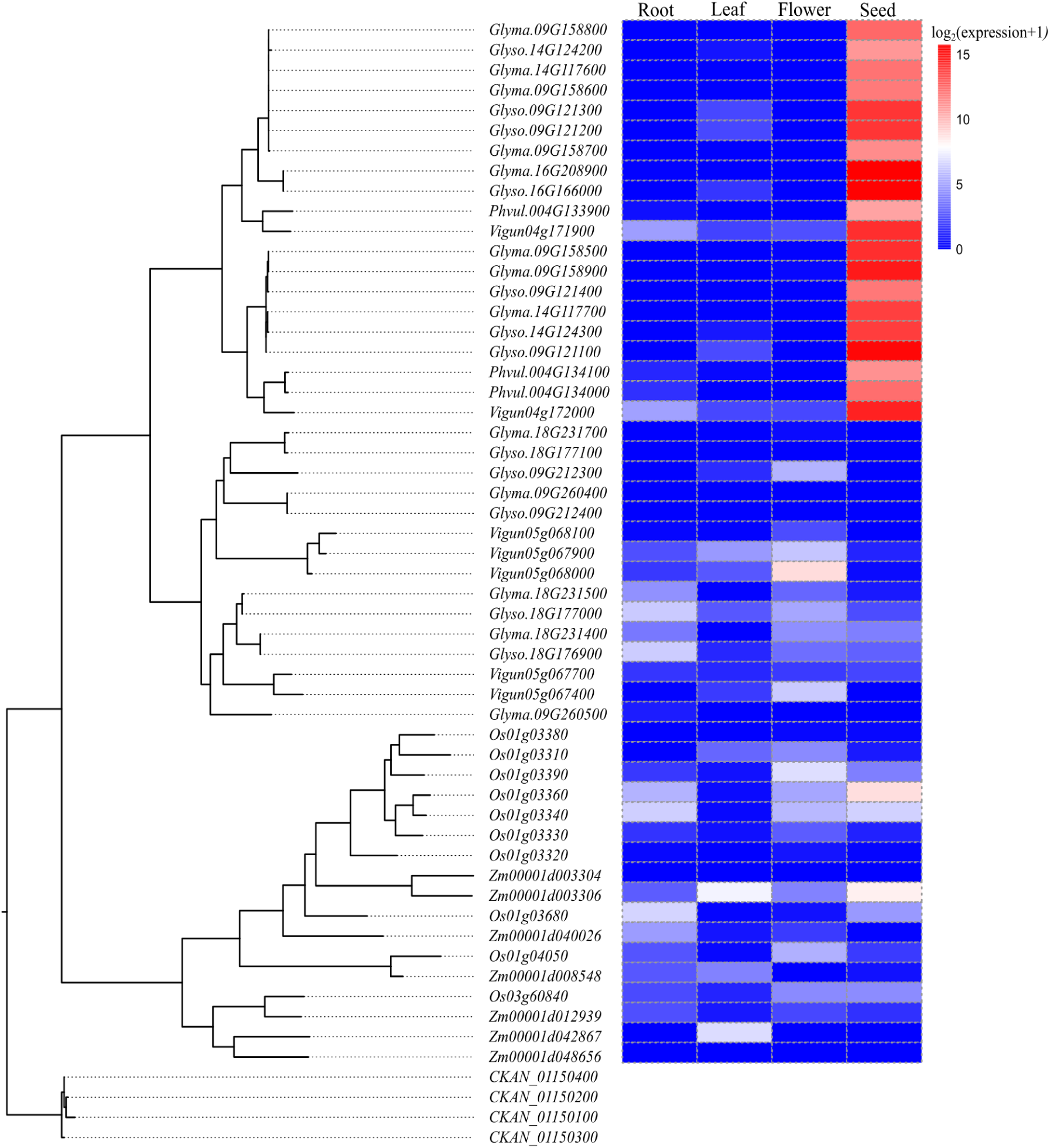
Phylogenetic tree and tissue-wide expression of BBI genes in four *Fabaceae* species (*Glycine max, Glycine soja, Vigna unguiculata, Phaseolus vulgaris*) and two *Poaceae* species (*Oryza sativa* and *Zea mays*). The Lauraceae species *Cinnamomum kanehirae* was used as an outgroup to root the phylogeny. Four tissues -root, leaf, flower, and seed-were used to investigate the expression patterns of BBIs.

### Soybean BBIs have evolved into a seed-specific subfamily and an environment-responsive subfamily

We examined the expression patterns of soybean BBI genes over the course of seed development (Figure 3A). Interestingly, each of the eight genes in subclade 1 showed little to no expression in seeds at embryogenesis stages (globular, heart and cotyledon stages). Expression was only evident at maturation stages, starting at early maturation stage, peaking at the mid-maturation stage, then gradually reducing to a low level at the dry seed stage. The five soybean BBI genes in subclade 2 exhibited no or very low expression in seeds at any seed development stage or leaf, root, stem and flower bud, raising the possibility that these genes are either pseudogenes or their transcriptional levels are regulated by environmental stresses. To determine whether any soybean BBI genes transcriptionally respond to environmental stresses, we investigated the expression pattern of each soybean BBI gene in response to 13 groups of environmental stresses including biotic stresses (e.g., fungal infection, virus infection), abiotic stresses (e.g., drought stress, salt stress) and combinatorial stimuli (e.g., nodulation and water restriction). BBI genes in subclade 1 did not transcriptionally respond to any examined abiotic and biotic treatments while all BBI genes in subclade 2 transcriptionally responded to abiotic and/or biotic stresses. However, each BBI gene in subclade 2 responded to a unique set of biotic and abiotic stresses (Figure 3B and Table S4). Each treatment regulated the expression of at least one *BBI* gene in Subclade 2. Specifically, expression of *GmBBI07* was regulated by *Rhizobial* nodulation and iron stress, while the expressions of *GmBBI11*, *GmBBI12* and *GmBBI13* changed significantly in response to multiple abiotic and biotic stresses. These three genes respond transcriptionally to ethylene treatment, abiotic stress signals including salt and drought stress, and biotic stresses like insect and fungal infections (Table S4 and Figure 3B). In addition, the expressions of *GmBBI11* and *GmBBI12* were regulated by treatments associated with circadian rhythm, common cutworm (CCW) and soybean cyst nematode (SCN).

**Figure 3.**
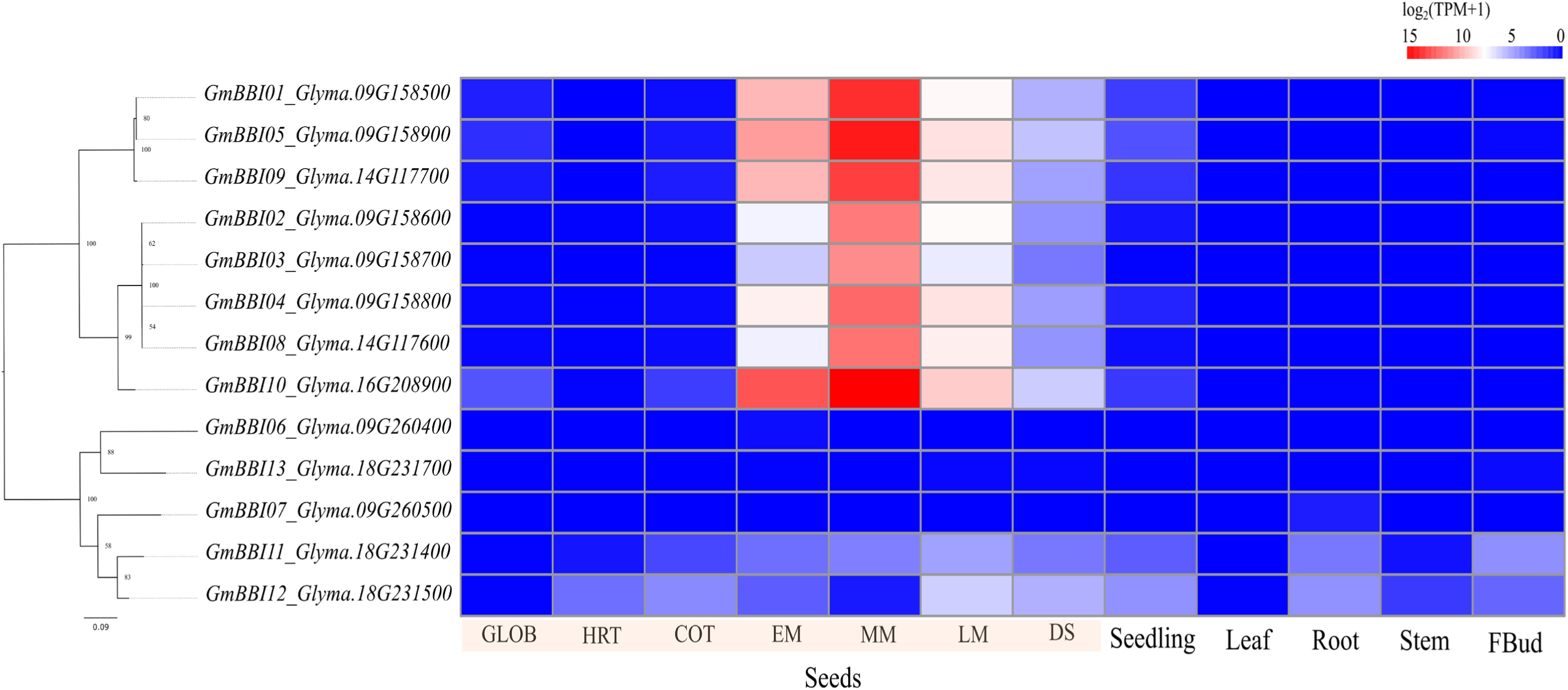
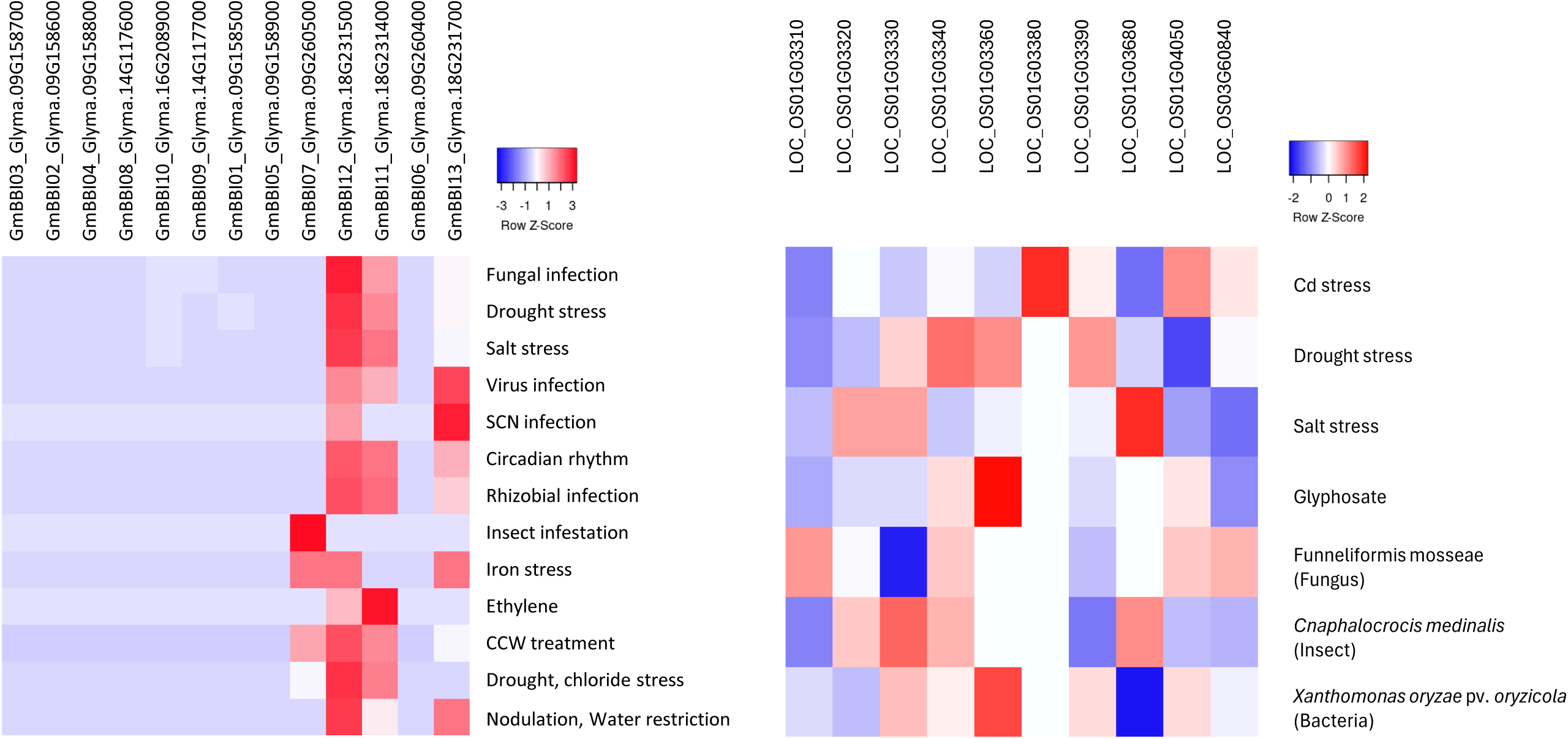
Heatmap of BBI gene expression across tissues and environmental signals. (A) Expression levels of BBI genes across various tissues are shown. These include seed developmental and maturation stages—GLOB (globular), HRT (heart), COT (cotyledon), EM (early maturation), MM (middle maturation), LM (late maturation), and DS (dry seed)—as well as tissues such as leaf, root, stem, and bud. The phylogenetic tree of 13 BBI genes was constructed depending on their amino acid sequences and displayed on the left. The heatmap shows the frequency of Treatment/Control (T/C) pairs where BBI genes in soybean (B) and rice (C) are detected for specific categories, including abiotic stress, biotic stress, phytohormone treatment, and other conditions.

Thus, there is a clear distinction in the transcriptional expression patterns of *BBI* genes in the two subfamilies (Figure 3 and Table S4). It is likely that the functions of the two BBI subfamilies are respectively associated with seed maturation and plant response to environmental stresses. We therefore refer to the two subfamilies as “seed-specific” and “environment-responsive” subfamilies.

BBIs are very ancient and emerged prior to the rise of angiosperms. *BBIs* in cereals have little or no expression in seeds (Figure 2) but have been shown to be transcriptionally regulated by abiotic and biotic stresses, similar to the clade 2 soybean BBIs (Shan *et al*., 2008; Xie *et al*., 2021; Zhang *et al*., 2024). Thus, legume subclade 1 BBIs may have acquired seed-specific expression and specific functions associated with seed maturation after divergence of subclass 1 and 2 while subclade 2 likely retained its ancestral stress-response function following the evolutionary divergence of the two subclades. Notably, rice BBIs also respond to environmental signals (Figure 3C). For example, LOC_Os01g03680 is upregulated under salt stress but downregulated during infection with the bacterial pathogen *Xanthomonas oryzae* pv. *oryzicola* (*Xoc*). Conversely, LOC_Os01g03360 is upregulated by Xoc, glyphosate, and drought stresses. These expression patterns highlight the conserved role of rice BBIs and legume subclade 2 in stress adaptation and defense mechanisms.

Neither seed maturation nor plant response to environmental stresses are essential for plant development and growth. BBI genes are unlikely to play essential functions in plant growth and development. Instead, the conservation of the environment-responsive BBI subfamily and acquisition of a newer seed-specific subfamily in legumes may contribute to fitness. This also explains why many species have lost their BBI genes without excessive adverse consequences to normal growth and development.

### Molecular mechanism of soybean BBI gene family expansion

Among the examined legume and cereal species, cultivated soybeans have the largest BBI gene family. Soybean BBI genes other than GmBBI10 (Glyma.16G208900) cluster on four distinct genomic regions of Chr09, 14, and 18 (Figure 4A and Table S5). Each cluster contains two-five tandem duplicated genes. Soybean, an allotetraploid species, has undergone two major whole genome duplication (WGD) events approximately 59 and 13 million years ago (Roulin *et al*., 2013). Macrosynteny analysis revealed homologous synteny blocks between chromosomes 09 and 16, and 09 and 18. These blocks contain homologous BBI genes, GmBBI01 (Glyma.09G158500) and GmBBI10 (Glyma.16G208900), and GmBBI06 (Glyma.09G260400) and GmBBI11 (Glyma.18G231400), attributable to the whole genome duplication events (Figure 3A). These two WGD-duplicated gene pairs respectively belong to the seed-specific and responsive soybean BBI subfamilies (Figure 2, Figure 4A, Table S5).

**Figure 4.**
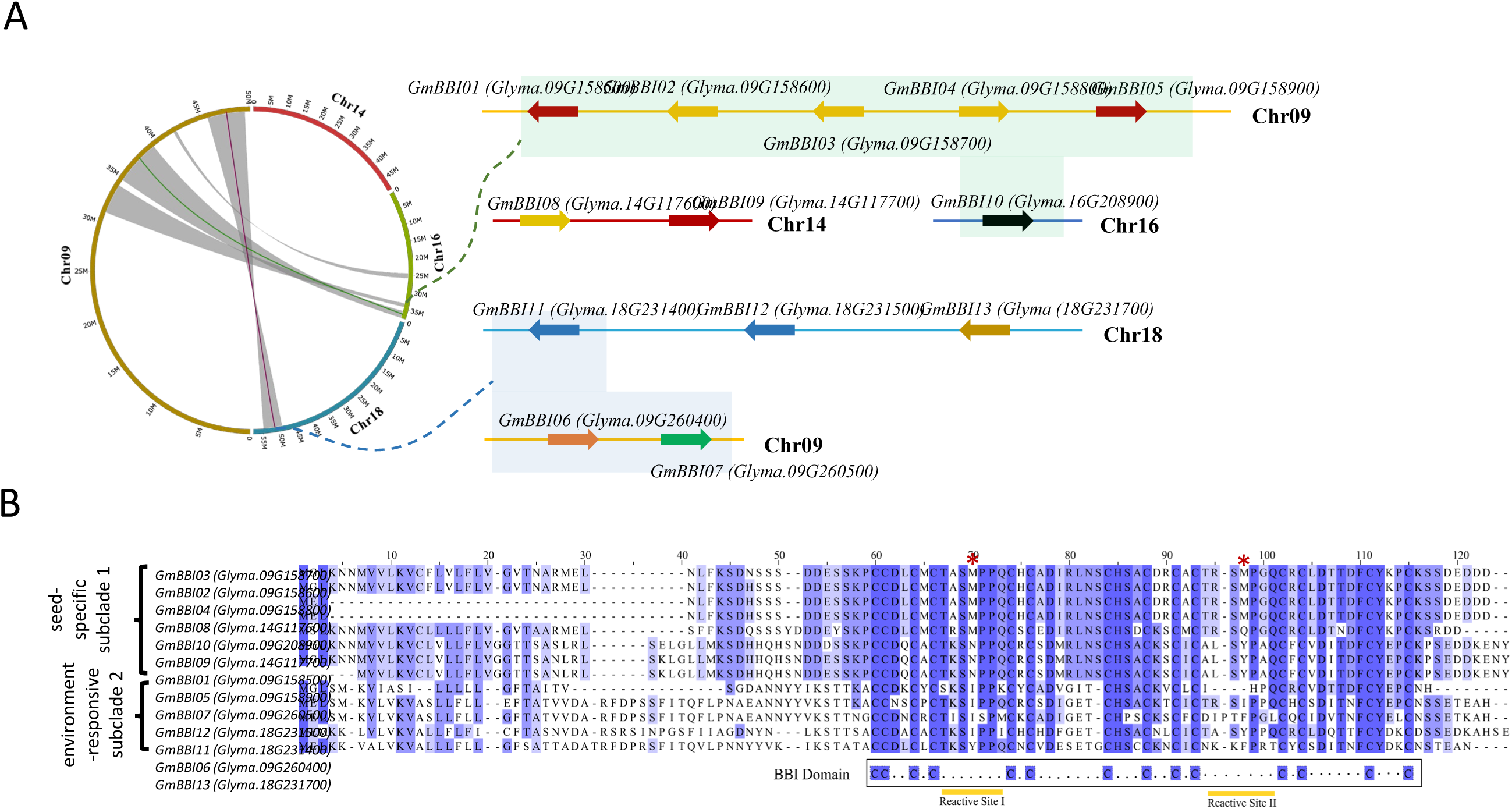
Gene duplication has expanded the soybean BBI gene family. **(A)** Macrosynteny and Microsynteny relationships of BBI genes across soybean chromosomes 09, 14, 16, and 18. The circos plot provides a genome-wide perspective, showcasing synteny between these chromosomes harboring BBI genes. Detailed views of microsyntenic blocks, specifically highlighted in light green and magenta boxes, focus on regions encompassing BBI genes. Additionally, the gene duplication is highlighted by coloring the duplicated genes with the same color for underscoring both tandem and segmental duplications. **(B)** Multiple sequence alignment of protein sequences of the BBI gene family members in soybean.

In addition to whole genome duplication, we observed that segmental and tandem duplications also contributed to BBI gene family expansion. Using DupGen_finder (Qiao *et al*., 2019) and a protein and coding sequence similarity threshold of 75%, we identified three pairs of tandem duplications (GmBBI03/GmBBI02, GmBBI04/GmBBI03, and GmBBI11/GmBBI12) along with six pairs of segmental duplications (GmBBI09/GmBBI10, GmBBI04/GmBBI08, GmBBI01/GmBBI05, GmBBI05/GmBBI09, GmBBI02/GmBBI04, and GmBBI03/GmBBI08) (Figure 4A and Table S5). All duplicated genes showed either seed-specific or environment-responsive expression patterns. Only one environment-responsive BBIs appears to have undergone tandem duplication, while seven seed-specific BBIs underwent tandem duplication. The low Ks values in the duplicated pairs of GmBBI02/GmBBI03, GmBBI03/GmBBI04, and GmBBI03/GmBBI08 suggest that they underwent duplication relatively recently (Table S5). Additionally, the BBIs in WGD pairs share only 42% (GmBBI01/GmBBI10) and 52% (GmBBI06/GmBBI11) protein sequence identities, while the tandem and segmental duplicated pairs share more than 75% protein identities (Table S5), consistent with their phylogeny (Figure 2). This implies that the tandem and segmental duplication events occurred after the whole genome duplication. Taken together, these findings suggest that segmental and tandem duplications are the main driving forces behind a recent expansion of the soybean BBI gene family, leading to a higher number of seed-specific BBI genes in particular.

### Preferential preservation of functional structures and activities of soybean BBI proteins

Multiple sequence alignment of Bowman-Birk inhibitor (BBI) peptides shows strong conservation of the BBI domains within and between two soybean BBI subfamilies (Figure 4B). In contrast, the N-terminal regions of the 13 soybean BBI open reading frames are more diverse in sequence. In addition, BBI sequences within each subfamily are more similar to each other than to the BBIs in the other subfamily.

Using SignalP 6.0 (Teufel *et al*., 2022), we predicted that at least 11 of the 13 BBIs possess an endoplasmic reticulum (ER) localization signal within the first 24th, 25th, or 27th amino acid (Figure S1A, C, D). However, two segmentally duplicated genes, *GmBBI04 and GmBBI08*, lack this ER signal (Figure S1B). AlphaFold structural analysis of the putative processed soybean BBIs (with ER signal peptides removed) showed that they all contain two inhibitory protease-binding loops, with no notable structural differences between seed-specific (Figure 5A) and environmentally responsive BBIs (Figure 5B). The 12–15 residue sequence segments that precede the conserved BBI domains in seed-specific BBIs exhibit greater sequence conservation than those in environmentally responsive BBIs (Figure 4B). However, these regions had less conserved and likely more flexible structures (Figure S2A-B), potentially leading to specialized functions for seed maturation and storage and responses to environmental stress.

**Figure 5.**
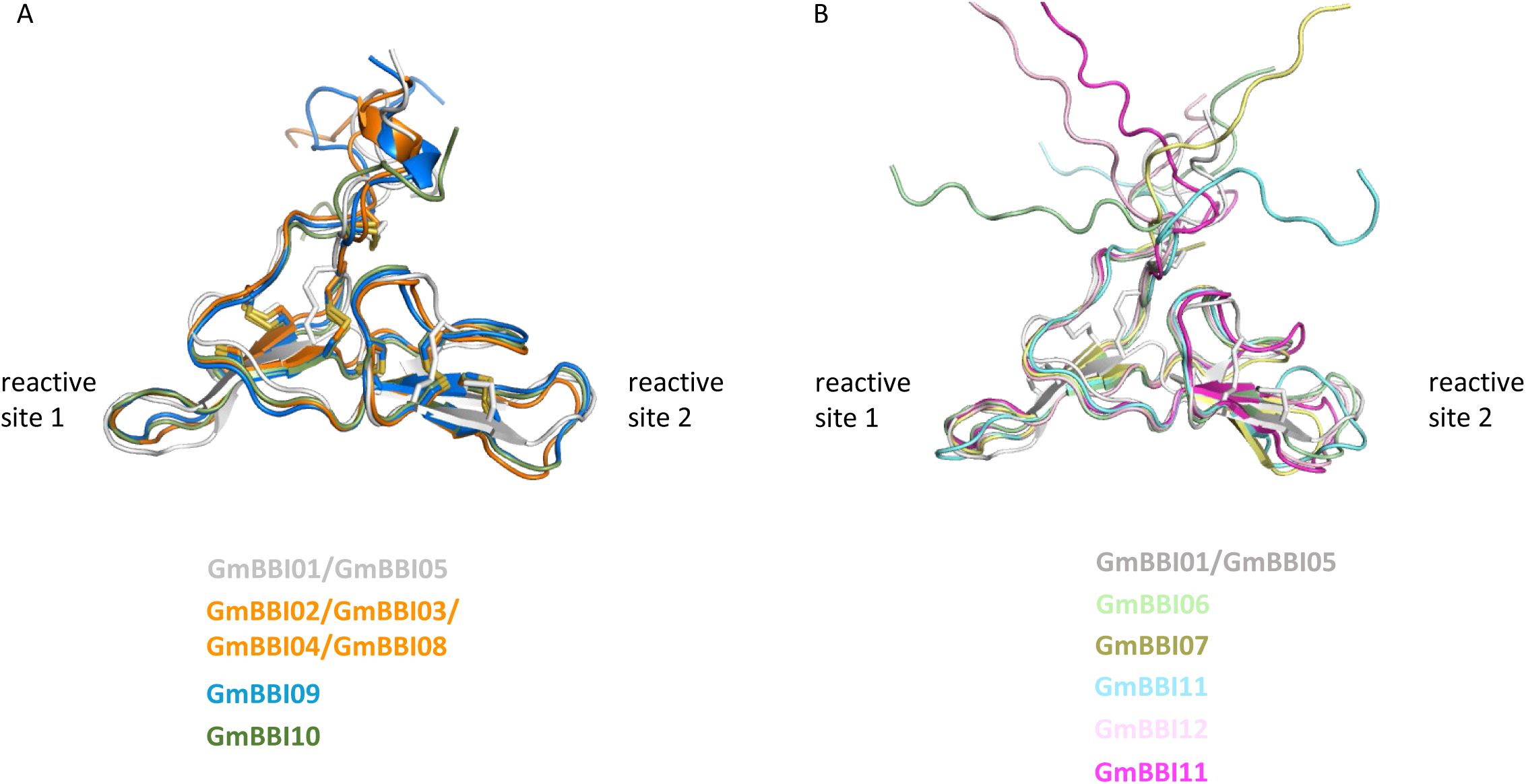
Structural alignment of the top AlphaFold models of soybean BBIs’ processed models. (A) Structural alignment between the averaged NMR structure of GmBBI01/GmBBI05 (PDB: 1BBI) and six other soybean BBIs in the seed-specific subfamily. (B) The alignment between the structure of GmBBI01/GmBBI05 and five soybean BBIs in the non-seed-specific and environmentally responsive subfamily. Note that the sequences of fully processed GmBBI01 and GmBBI05 are identical.

To explore protease binding, we used AlphaFold2 to model interactions between the soybean BBIs and trypsin or chymotrypsin, analyzing the five highest-scoring models for each complex. All soybean BBIs were predicted to bind trypsin in a similar manner as the previously experimentally determined GmBBI1:trypsin complex ((Koepke *et al*., 2000; Tornøe *et al*., 2017) (Figure S3). For seed-specific BBIs GmBBI01, GmBBI05 (Figure S3A), GmBBI09 (Figure S3C), and GmBBI10 (Figure S3D), as well as environmentally responsive BBIs GmBBI06 (Figure S3E), GmBBI12 (Figure S3H), and GmBBI13 (Figure S3I), the top models included binding via either reactive site 1 (blue) or 2 (yellow), resembling the binding modes experimentally observed for GmBBI1. However, models for seed-specific BBIs GmBBI02, GmBBI03, GmBBI04, and GmBBI08 exclusively showed binding via reactive site 2 (Figure S3B), while environmentally responsive BBIs GmBBI07 (Figure S3F) and GmBBI11 (Figure S3G) consistently bound via reactive site 1.

Similar trends were observed for chymotrypsin binding (Figure S4). All eight seed-specific BBIs Figure S4A-D) and three environmentally responsive BBIs, GmBBI06, 12, 13 (Figure 45E, 4H and 4I) showed flexibility in binding via either reactive site 1 or 2. In contrast, environmentally responsive BBIs, GmBBI07 and GmBBI11, favored reactive site 1 for chymotrypsin (Figure S4F, G). These in silico predictions of binding preferences and inhibitory activity require further experimental confirmation.

### Development of soybean containing reduced BBI activities by editing seed-specific BBI subfamilies

BBIs in soybean seeds have negative impacts on nutritional value of soybean for feed and food. To develop soybeans with reduced BBI activity in seeds and assess the role of seed-specific BBIs in soybean growth and development, we applied CRISPR-Cas9 technology to knock out the seed-specific genes. Having transformed soybean (WM82) with a vector containing four gRNAs targeted to the seed-specific genes (Figure S5A and S5B), we detected mutations in each of the eight specific BBI genes in transformed soybean (Figure S6). The mutations include DNA deletions as well as combinations of deletions and insertions. They introduce premature stop codons, cause reading frame shifts, and disrupt the conserved BBI functional domain (Figure S7A–C), which are expected to inactive BBI activity (Table S6).

We assessed TIA and CIA in T2 seeds of five edited lines, each containing one or more of the three homozygous BBI edited alleles of *GmBBI01 (GmBBI01-a), GmBBI08 (GmBBI08-a and -b), and GmBBI10 (GmBBI10-a and -b)* and heterozygous alleles for the remaining *BBIs* (Table S6). For the non-edited line WM82, we measured a TIA of 48.5 mg trypsin inhibited/g sample and CIA of 37.9 mg chymotrypsin inhibited/g sample. In the edited plants, we observed a drastic reduction in both TIA (by 69.6%-73.7%) and CIA (by 76.4 – 79.4%) (Figure 6A). We assessed the accumulated levels of BBI protein in the edited plants by SDS-PAGE. CRISPR-edited mutant seed extracts did not show any BBI protein band while non-edited soybean (wild-type) seed extracts displayed a BBI protein band at approximately 9.5 kDa (Figure 6B), confirming that BBI protein accumulation is significantly reduced in the edited seeds.

**Figure 6.**
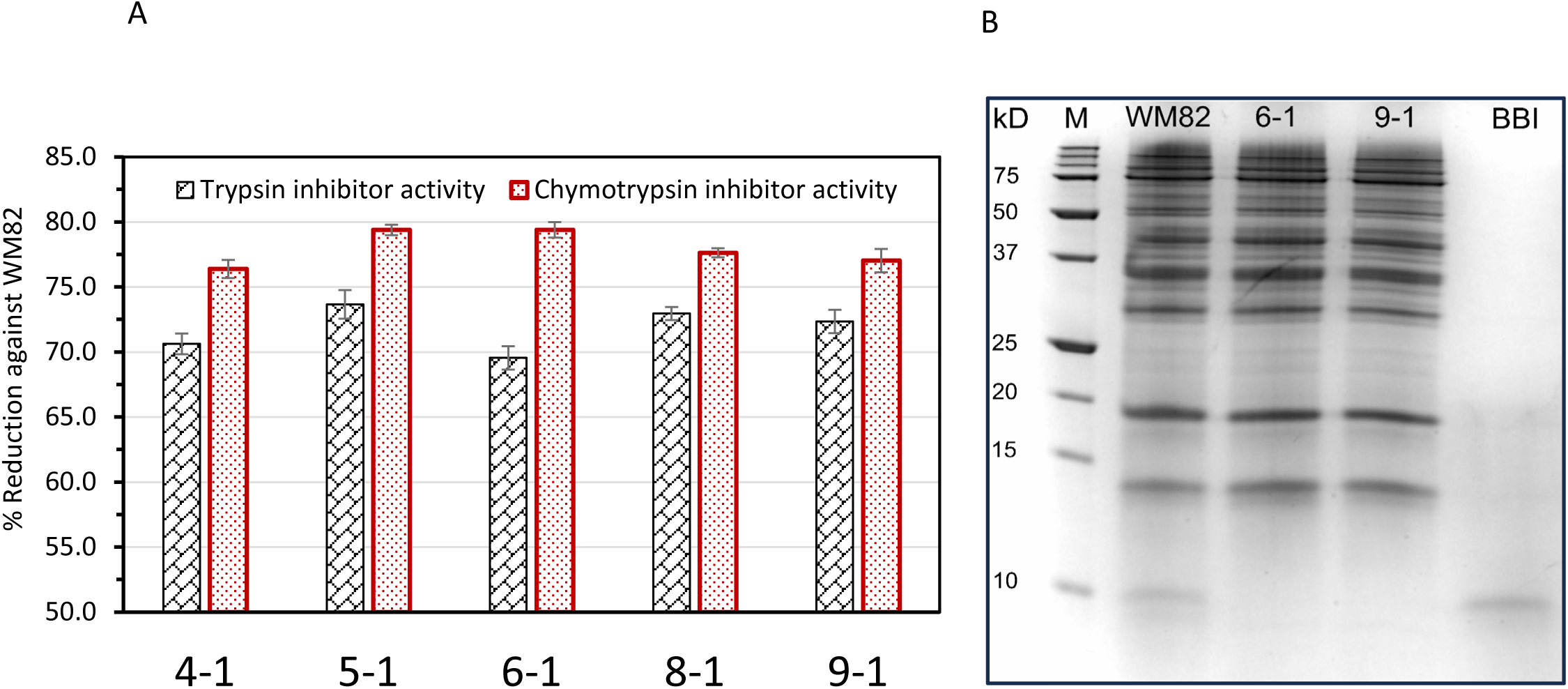
BBI content decreases dramatically in the gene-edited seeds compared to WM82. **(A)** Percent reduction in trypsin inhibitor activity (TIA) and chymotrypsin inhibitor activity (CIA) in T2 seeds of the CRISPR-Cas9 edited mutants against Williams 82 (wild-type). **(B)** Tricine SDS-PAGE of two mutant soybean seeds (T2), Williams 82 (WM82), and purified BBI. Lane M represents protein markers with descending MWs. Seed samples were loaded based on the same protein amount for each lane.

### No obvious phenotypic change in edited soybean with reduced BBI activity

We measured phenotypes of major traits associated with agronomic performance in the BBI-edited lines. We did not observe any significant changes in 100-seed weight, yield per plant, maturity, and plant height (Figure 7). The 100-seed weight and yield per plant ranged from 20.61g to 20.81g and 60.91g to 63.67g, respectively (Figure 7A). The gene-edited plants and WM82 matured in 109 to 114 days, with plant heights of 82 to 85 cm (Figure 7B). Furthermore, all BBI gene-edited seeds exhibited a 100% germination rate when cultivated under greenhouse conditions (Figure 7B). All the gene-edited plants appear normal visually, compared to the control WM82 (Figure 7C).

**Figure 7.**
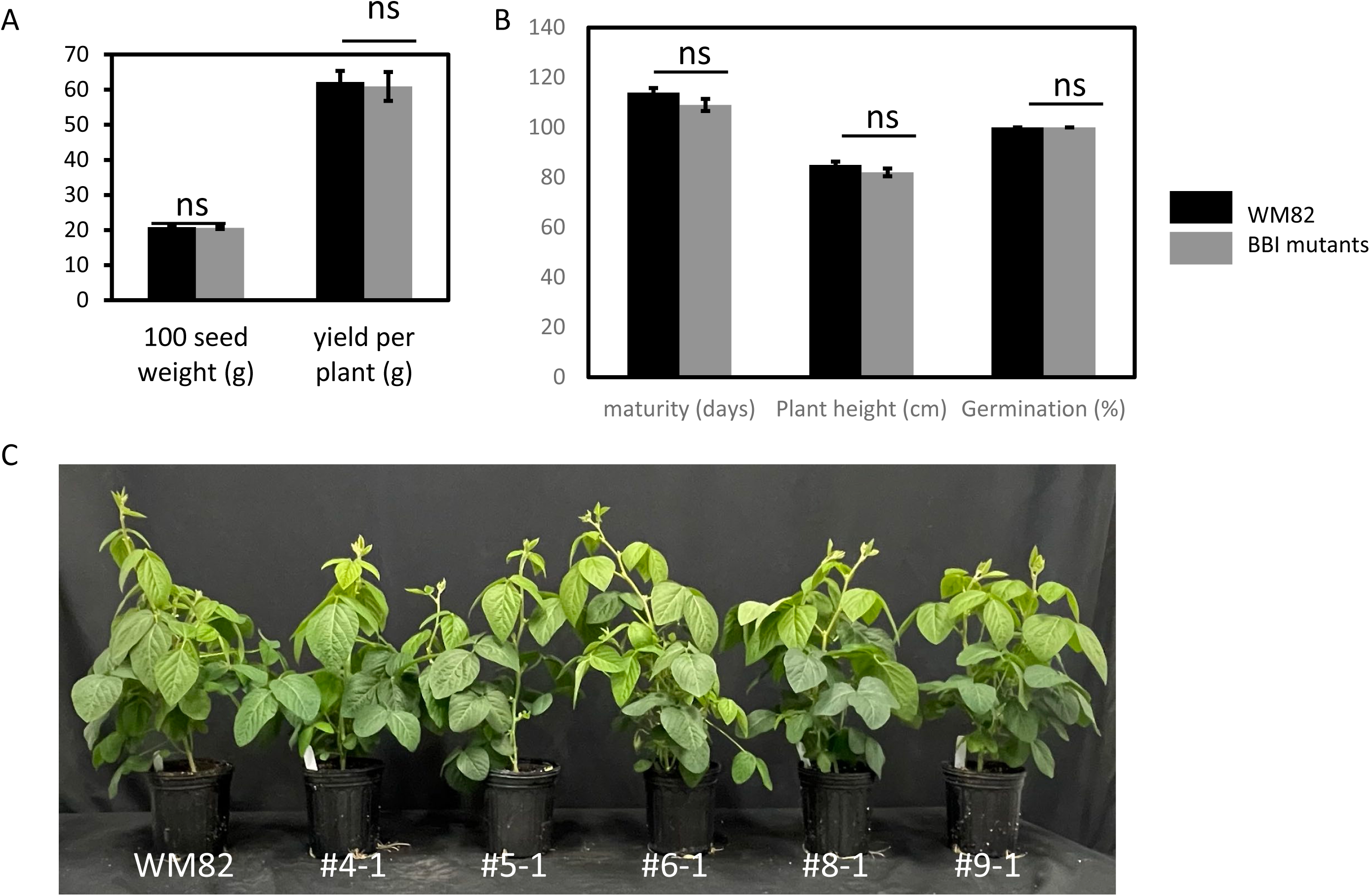
Agronomic traits of CRISPR/Cas9-mediated genome-edited seeds do not significantly alter compared to WM82. (A) displayed 100-seed weight (g) and yield per plant (g), while (B) exhibited maturity (days), plant height (cm), and germination rate (%). (C) The phenotypes of WM82 and gene-edited plants. Statistical significance is determined based on a p-value threshold of 0.05.

Next, we measured additional quality parameters related to seed composition. These included contents of protein, oil, simple sugars (sucrose, fructose, and glucose). Compared to the control seeds, no significant alteration was observed in all these compositional attributes (Table 2).

**Table 2.** Seed compositions of CRISPR/Cas9-mediated genome editing seeds, in comparison to non-edited William 82 (wild-type) 1,2.

| Sample ID<sup>3</sup> | Protein (%) | Oil (%) | Cysteine (%) | Methionine (%) | Sucrose (%) | Fructose (%) | Glucose (%) |
| --- | --- | --- | --- | --- | --- | --- | --- |
| Non-edited Wm 82 | 39.82 ± 1.55a | 19.58 ± 0.69a | 0.59 ± 0.04a | 0.53 ± 0.02a | 5.73 ± 0.48a | 0.70 ± 0.05a | 0.65 ± 0.01a |
| #4-1T | 39.51 ± 0.74a | 19.77 ± 0.76a | 0.61 ± 0.03a | 0.53 ± 0.03a | 5.69 ± 0.76a | 0.67 ± 0.08a | 0.65 ± 0.02a |
| #5-1T | 40.02 ± 1.21a | 19.67 ± 0.53a | 0.61 ± 0.03a | 0.52 ± 0.05a | 5.22 ± 0.65a | 0.72 ± 0.08a | 0.65 ± 0.01a |
| #6-1T | 39.50 ± 1.70a | 19.92 ± 1.14a | 0.59 ± 0.04a | 0.53 ± 0.05a | 5.73 ± 0.84a | 0.62 ± 0.09a | 0.65 ± 0.02a |
| #8-1T | 38.92 ± 1.71a | 20.02 ± 0.41a | 0.59 ± 0.03a | 0.51 ± 0.03a | 5.73 ± 1.03a | 0.65 ± 0.07a | 0.64 ± 0.01a |
| #9-1T | 38.75 ± 0.33a | 20.04 ± 0.39a | 0.57 ± 0.02a | 0.51 ± 0.03a | 5.63 ± 1.03a | 0.68 ± 0.04a | 0.64 ± 0.01a |

<sup>1</sup> The seed composition parameters are expressed as percent dry weight basis.
<sup>2</sup> Means in each column with different letters denote statistical significance ( $p < 0.05$ ).
<sup>3</sup> "T" represents transgenic lines.

BBIs contain seven disulfide bonds formed from 14 cysteine residues. Given the high relative density of cysteines, BBIs have been considered as storage proteins for sulfur and/or a rich source of sulfur-containing amino acids that contribute to soybean nutritional value. We did not observe statistically significant differences between seeds of WM82 and mutants in overall content of cysteine and methionine (Table 2). Thus, significant reduction of BBIs does not appear to affect soybean seed quality and agronomic traits, critical factors for the development of commercially viable low trypsin inhibitor soybean. Our results also support the notion that seed-specific BBIs do not play an essential function in plant growth and development.

## Discussion

### One Head or Two: Distinct Structures of BBIs in Cereals and Legumes

Bowman-Birk inhibitors (BBIs) are serine protease inhibitors with multiple conserved disulfide bonds, first identified in soybean (Bowman, 1946). Legume BBIs possess two reactive sites capable of simultaneously inhibiting trypsin and chymotrypsin, forming a “double-headed” structure (Birk, 1985). In contrast, cereal BBIs typically lack critical cysteine residues and retain only a single inhibitory loop, often unable to inhibit chymotrypsin effectively (Park et al., 2004; Qi et al., 2005). Our phylogenetic analysis supports this divergence, with legume and cereal BBIs forming two distinct lineages (Figure 1).

All 13 soybean BBI genes contain dual reactive sites (Figure 4B), and structural modeling suggests that at least 11 can simultaneously bind trypsin and chymotrypsin (Figure 5, Figure S3, Figure S4). Prior studies confirmed that GmBBI01 can bind both enzymes via its two sites (Koepke et al., 2000). We also observed that in GmBBI07 and GmBBI11, site 1 may functionally resemble the canonical site 2, indicating functional divergence between the two homologous loops.

Structural and sequence homologies across plant species indicate a common ancestral BBI, with the double-headed structure in legumes likely arising from an ancient gene duplication before angiosperm divergence. Selaginella moellendorffii, a lycopod predating angiosperms by 200–230 million years, possesses a double-headed BBI, supporting this hypothesis (James et al., 2017), suggesting domain duplication occurred before angiosperm emergence. Soybean BBIs retain the double-headed structure, likely enhancing fitness under natural conditions, while cereal BBIs have lost one head, reflecting greater structural variability in monocots (Qi et al., 2005).

### Molecular Mechanism of soybean BBI gene family expansion

Compared to other legumes such as common bean, cowpea, and peanut, soybean contains the highest number of BBI genes within its genome (Figure 1 and Table 1). Our synteny analysis revealed that two whole genome duplication events - on Chromosomes 9 (GmBBI01) and 16 (GmBBI10), and Chromosomes 9 (GmBBI06) and 18 (GmBBI11) - expanded the soybean BBI gene family approximately 13 million years ago (Figure 4A and Table S5). Three tandem duplications on Chromosomes 9 and 18, and six segmental duplications on Chromosomes 9 and 14 have further expanded the family (Figure 4A and Table S5). While the two duplicated pairs in WGD blocks have 42% and 52% protein identities, the tandem and segmental duplicated pairs share more than 75% protein identities. For instance, GmBBI02 and GmBBI03, a tandem duplicated pair, have identical amino acid sequences. GmBBI04 and GmBBI08, a segmental duplicated pair, both have a deletion in their N-terminus compared to other BBIs. Compared to WGD pairs, tandem and segmental duplicated pairs have much lower Ks values (Table S5). Thus, diverse molecular mechanisms of gene duplication led to the expansion of the soybean BBI gene family, with tandem and segmental duplications representing the predominant driving forces for its recent expansion.

### Legume BBIs evolved into two divergent subfamilies with distinct and conserved seed-specific and environment-responsive expression patterns

Beyond gene duplication and protein structure comparisons, our study presents new insights into molecular evolution, regulation and functions of BBIs. We conducted a gene expression analysis of BBI genes in two cereals (maize and rice), four legumes (cultivated soybean, wild soybean, common bean, and cowpea) (Figure 2). Cereal BBIs were expressed only at low levels in the examined tissues. However, the two divergent legume BBI subfamilies showed distinct and conserved expression patterns (Figure 2). The expression profiles of BBIs from these seven species are consistent with their phylogeny (Figure 1). The coevolution of BBI sequences and gene expression patterns and highly divergent expression patterns of the two subfamilies suggest that they have distinct biological functions associated with seed development and response to environmental changes respectively. Interestingly, no obvious duplication signature is present across the two subfamilies of soybean BBIs, implying that the expression patterns and functional characteristics were determined before BBI family expansion.

From the perspective of protein sequence analysis, the BBI domain in the C-terminal portions and the ER peptide signals at the N-terminal portions of BBI genes from the two subfamilies are both largely conserved, suggesting that they are subject to functional constraints (Figure 4B and Figure S1). This structural variability, in contrast to the conservation of the BBI domains, suggests that evolutionary selective pressure has acted primarily to maintain the functional integrity of the BBI domain.

Our detailed analysis of BBI expression patterns supports the distinct functions of the two subfamilies of BBI genes in soybeans. Our analysis reveals the conserved and specific expression of the eight seed-specific BBIs in seeds of all examined legume species, while members of the other BBI subfamily members are expressed in response to abiotic and biotic stresses rather than in tissue-specific patterns (Figure 3 and Table S4). These differential expression patterns further support the two soybean BBI subfamilies have followed independent evolutionary pathways. In addition, each gene in the environment-responsive subfamily evolved its own pattern of transcriptional response to environmental changes, consistent with proposed protective functions against diverse abiotic and biotic stresses in nature. Gene expression changes quickly if a gene is not subject to functional constraints. Preserving the conserved and distinct expression of each subfamily suggest that they are subject to functional constraints (Shakhnovich and Koonin, 2006)

Seed development is divided into embryogenesis and maturation. A fertilized cell divides and differentiates into an embryo during seed embryogenesis while seeds further enlarge, produce storage reserves, desiccate and develop into dormancy during seed maturation. However, seed maturation is considered as a non-essential and adaptive biological process since an embryo from embryogenesis can geminate and finish its entire life cycle without seed maturation. In the present study, all seed-specific BBI genes were primarily expressed at maturation stages, not at embryogenesis stages (Figure 3A). The conserved and specific expression of eight seed-specific BBIs at seed maturation stage, but not in embryogenesis, suggests that the seed-specific genes are unlikely to have essential functions in plant growth and development, but rather act to increase plant fitness by natural selection. We observed no obvious phenotypic changes following significant reduction of BBI activity. The absence of BBI genes in species such as Arabidopsis, and the fact that rice and maize *BBI* genes show little or no seed-specific expression, further supports this conclusion.

Several plant protease inhibitors have been reported to play roles in regulation of endogenous protease activity associated with seed development, storage protein accumulation, seed germination and defense, which are important for plant fitness (Clemente *et al*., 2019). BBI has been suggested to play a role in protection against birds and herbivores, and has been termed a “biopesticide” due to its ability to inhibit trypsin activity and reduce protein digestion efficiency in animal gut (Rodriguez-Sifuentes *et al*., 2020; Sultana *et al*., 2023; Wang *et al*., 2015; Xie *et al*., 2021). BBI has been engineered to develop insect-resistant transgenic crops, in which overexpression of a cowpea BBI gene confers resistance to Coleopteran and Lepidopteran insects (Bi *et al*., 2006; Hilder *et al*., 1987; Xu *et al*., 1996). Seed-specific BBIs may also function as cysteine-enriched seed storage proteins and play a protective role for protein-rich seeds against herbivores.

In contrast, the other BBI sub-family genes respond highly to various environmental stimuli, including SCN, viruses, rhizobia, fungi, common cutworms, ethylene, salt, and drought (Figure 3B and Table S4). Their expression patterns in response to environmental stresses are highly diverse. The expression of rice BBI genes was also found to be responsive to environmental conditions (Figure 3C). Roles for BBIs in environmental responses has been previously reported for both cereals and legumes (Drame *et al*., 2013; Juliana *et al*., 2018; Malefo *et al*., 2020; Sari *et al*., 2019; Shan *et al*., 2008; Zhang *et al*., 2024). The subfamily is ancient and evolutionarily conserved across plant lineages and plays roles in response to diverse abiotic and biotic stimuli, enhancing the plant’s overall fitness (Figure 3). In contrast, the legume seed-specific BBIs likely underwent sub-functionalization in gene expression, diverging from their environmentally responsive counterparts. This suggests that the seed-specific subfamily arose more recently and evolved to a new biological function related to the seeds, increasing plant fitness. For example, seed-specific BBIs could enhance seed survival in harsh winter or defend against herbivores. It will be important to examine how knockouts of individual BBI genes affect resistance of soybean to the range of applicable environmental stresses.

### Implications for soybean and legume nutrition improvement and future research

Legume seeds are one of the most important sources of proteins for feed and food. However, high accumulation of BBIs in seeds reduce the digestibility of seed proteins and contributes to stomach problems, due to inhibition of gastrointestinal digestive protease activities in animals. Many studies have been performed to understand BBI functions due to their significance in agriculture of soybean and other legumes. Due to lack of comprehensive understanding of the molecular properties and functions of the BBIs, we still confront the dilemma of designing effective strategies to reduce or eliminate BBI activity without negatively impacting plant growth and development or the quality of the harvested product. Our comprehensive evolutionary and molecular study indicates that BBIs in soybeans and legumes are encoded by a large gene family. BBIs in seeds are mainly produced by eight genes in a seed-specific subfamily. All of them are specifically expressed in seeds, but not in the other examined tissues or response to environmental stresses. Expression of the soybean seed-specific BBI subfamily was restricted to seeds during non-essential maturation stage, but not during essential embryogenesis over the course of soybean growth and development. Together with absence of BBI genes in many angiosperm species (James *et al*., 2017), the lack of expression of the seed-specific BBI subfamilies in the other tissues suggest that knockout of the seed-specific BBI genes unlikely affect plant growth and development. Our findings support the feasibility of developing commercially viable soybean that contains reduced or no BBI protein in seeds without affecting other major phenotypes. Therefore, we targeted seed-specific BBI genes by gene-editing. We reduced BBI activity by 70% in soybeans containing various combinations of homozygous and heterozygous edited seed-specific BBI genes (Figure 6A). At the same time, important agronomic traits, including plant height, maturity, 100-seed weight, and yield per plant, showed no significant differences between the non-edited and edited soybeans (Figure 7). Notably, we examined the germination rate of the edited soybean lines and did not observe any significant differences, even though BBI potentially play a role in hydrolyzing and mobilizing reserve proteins to support seed germination seed (Martinez *et al*., 2019).

Overall, the study shows that BBIs are encoded by a large and complex gene family that has undergone expansion and divergence. We propose that the seed-specific subfamily potentially plays roles in enhancing seed resilience to abiotic and biotic stresses such as protecting seeds from harsh environmental stresses and against anti-herbivory, while the stress-responsive subfamily is likely involved in defense against abiotic and biotic stresses, rather than in plant growth and development. Our findings point to a potential conflict between BBIs’ evolutionary roles in natural selection and their current use as feed and food in agriculture but also provide insights applicable to the development of low-BBI soybean cultivars that maintain agronomic performance. The expansion of the seed-specific BBI subfamily, preservation of seed-specific expression for the eight BBI genes at seed maturation, and inhibitory activity of protease in animals suggest that they function in protecting soybean seeds from herbivores and other harsh environmental stresses. Accumulation of BBIs in seeds is likely beneficial in natural evolution and potentially increases soybean fitness by protecting soybean seeds from herbivores and other stresses but is undesirable for the current use of soybean as food and feeds. This conflict is potentially resolvable as modern soybean agricultural practice provides controlled optimized growth conditions that decrease the effects of herbivores and increase seed viability. Since soybean seed-specific BBI genes are only expressed in maturing seeds, not developing seeds during embryogenesis, non-seed tissues or response to environmental stresses, significant reduction of seed-specific BBIs should not affect plant development or interaction of soybean with diverse abiotic and biotic stresses. Our future research will focus on evaluating the field performance of homozygous knockouts of all eight seed-specific BBI genes; the resulting findings will provide more affirmative knowledge about BBI roles and validate novel approaches and solutions to the challenge of soybean crop improvement.

## Author contributions

YQA conceived, led and participated in all aspects of the study. JH conceived, initiated and generated significant preliminary data and discoveries for the study. ZW screened and determined phenotypic data of edited plants and drafted the manuscript. LGL carried out phylogenetic analysis. NL conducted research on gene family expansion and expression patterns for different species. WS and NL performed gene expression data analysis for BBIs in response to internal and external signals. SM performed protein structural modeling and analysis, SP designed guide-RNAs for 8 BBI genes, RC and SMK constructed the vector and generated transformed soybean plants, KL carried out TIA and CIA assays and SDS-PAGE. All authors reviewed and approved the manuscript.

## Supporting information

BBI_BioRxiv_Figures S1-S7_Sub

## Acknowledgements

This work was supported by the United Soybean Board (**# 2332-203-0103 and 24-201-S-C-1-A**) to YQA and by USDA-ARS funded to YQA and KL. We would like to thank Drs. Dilip Shah and Mao Li for their scientific and managerial help, Dr. Wolfgang Goettel for scientific input and Mr. Rick Meyer and Mr. Mike Woolman for their technical support.

## Disclaimer Note

Names are necessary to report factually on available data; however, the USDA neither guarantees nor warrants the standard of the product, and the use of the name by USDA implies no approval of the product to the exclusion of others that may also be suitable. USDA is an equal opportunity provider and employer.

## Legends to Supplementary Figures

**Figure S1. The N-terminal amino acid sequences in 11 out of the 13 soybean BBIs function as endoplasmic reticulum signal peptides.** (A) GmBBI01, GmBBI02, GmBBI03, GmBBI05, GmBBI07, and GmBBI09 contain endoplasmic reticulum signal peptides within their first 25 amino acids. (B) GmBBI04 and GmBBI08 do not contain significant signal peptides. (C) GmBBI06, GmBBI11, GmBBI12, and GmBBI13 contain endoplasmic reticulum signal peptides within their first 27 amino acids. (D) GmBBI10 contains endoplasmic reticulum signal peptides within their first 24 amino acids.

**Figure S2. Modeling of pre-processed 13 soybean BBI proteins.** The top 5 AlphaFold models of the pre-processed amino acid sequences are superimposed and displayed for the two subfamilies of soybean BBIs: (A) seed-specific subfamily, and (B) environment-responsive subfamily.

**Figure S3. Structural comparative analysis of soybean BBIs complexed with bovine trypsin.** The PDB: 1d6r model (Koepke *et al*., 2000) complex between trypsin and the processed forms of two identical soybean BBI genes, GmBBI01 and GmBBI05, serves as the reference for other complexes involving trypsin and different soybean BBIs. Alphafold predicted the top 5 model complexes between trypsin and various soybean BBIs as follows: (B) GmBBI02/GmBBI03/GmBBI04/GmBBI08, (C) GmBBI09, (D) GmBBI10, (E) GmBBI06, (F) GmBBI07, (G) GmBBI11, (H) GmBBI12, and (I) GmBBI13.

**Figure S4. Structural comparative analysis of soybean BBIs complexed with chymotrypsin.** The PDB: 5j4q model (Tornøe *et al*., 2017) complex between chymotrypsin and the processed forms of two identical soybean BBI genes, GmBBI01 and GmBBI05, serves as the reference for other complexes involving chymotrypsin and different soybean BBIs. Alphafold predicted the top 5 model complexes between trypsin and various soybean BBIs as follows: (B) GmBBI02/GmBBI03/GmBBI04/GmBBI08, (C) GmBBI09, (D) GmBBI10, (E) GmBBI06, (F) GmBBI07, (G) GmBBI11, (H) GmBBI12, and (I) GmBBI13.

**Figure S5. Map of the binary vector used for CRISPR/Cas9 mediated gene editing on GmBBIs. (A)** The CRISPR/Cas9 construct harbors three necessary elements exhibited as below: the selection cassette consists of 35S promoter, spectinomycin resistant gene (soybean transformation selection marker), and 35S terminator; the Cas9 cassette consists of ubiquitin promoter, Cas9 gene, and ubiquitin terminator; four guide RNA (gRNA) cassettes. **(B)** The sequences of four sgRNAs and their target genes are shown here. Only GmBBI04 (Glyma14g117600) was targeted by one gRNA while other seven BBI genes were targeted by two gRNAs.

**Figure S6. Heterozygous edited genotypes of (A) GmBBI02 (Glyma.09g158500), (b) GmBBI03 (Glyma.09g158600), (C) GmBBI04 (Glyma.09g158700), (D) GmBBI05 (Glyma.09g158800), (E) GmBBI09_Glyma.14g117700.** The sequence in guiding RNA region is indicated.

**Figure S7. The CRISPR/Cas9-mediated genome editing approach successfully manipulates the target BBI genes.** (A) Mutated genotype of GmBBI01 (Glyma.09g158500) was identified in line 9-1T, where 31 bp were deleted between two gRNAs. The amino acid alignment indicates that this mutation causes a premature stop codon at 17^th^ amino acid. (B) Mutated genotype 1 of GmBBI08 (Glyma.14g117600) was identified in lines 4-1T and 8-1T, where 2 bp was deleted in the gRNA regions. Mutated genotype 2 of GmBBI08 was identified in lines 6-1T and 9-1T, where 4 bp were deleted in the gRNA region. The amino acid alignment indicates that mutation 1 causes a premature stop codon at 40^th^ amino acid while mutation 2 causes a premature stop codon at 39^th^ amino acid. (C) Mutated genotype 1 of GmBBI10 (Glyma.16G208900) was identified in lines 5-1T and 8-1T, where 48 bp were deleted between two gRNAs, respectively. Mutated genotype 2 of GmBBI10 was identified in lines 6-1T and 9-1T, where a loss of 55 bp was accompanied by an insertion of 23 bp between the two gRNAs. The amino acid alignment indicates that mutation 1 causes a truncation at N-terminus while the mutation 2 causes a premature stop codon at 14^th^ amino acid.

