## Supplementary material for "Divergence of Bowman-Birk Protease Inhibitor Family into seed-specific and environmentally responsive subfamilies in Legume and Soybean: Implication for Legume Seed Composition Improvement": BBI_BioRxiv_Figures S1-S7_Sub: BBI_BioRxiv_Figures S1-S7_Sub.pptx

### Slide 1
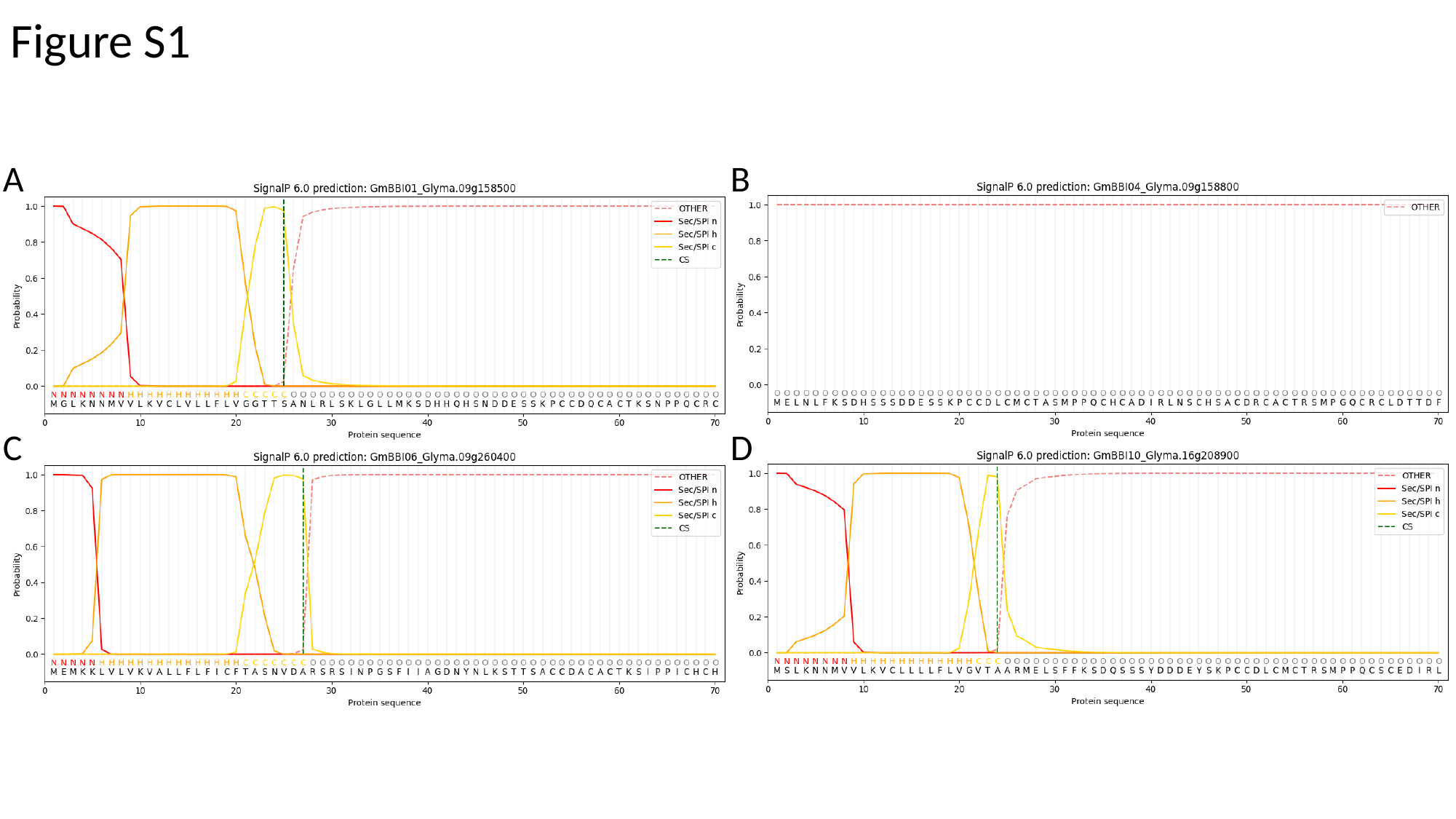

Figure S1
A
B
C
D

### Slide 2
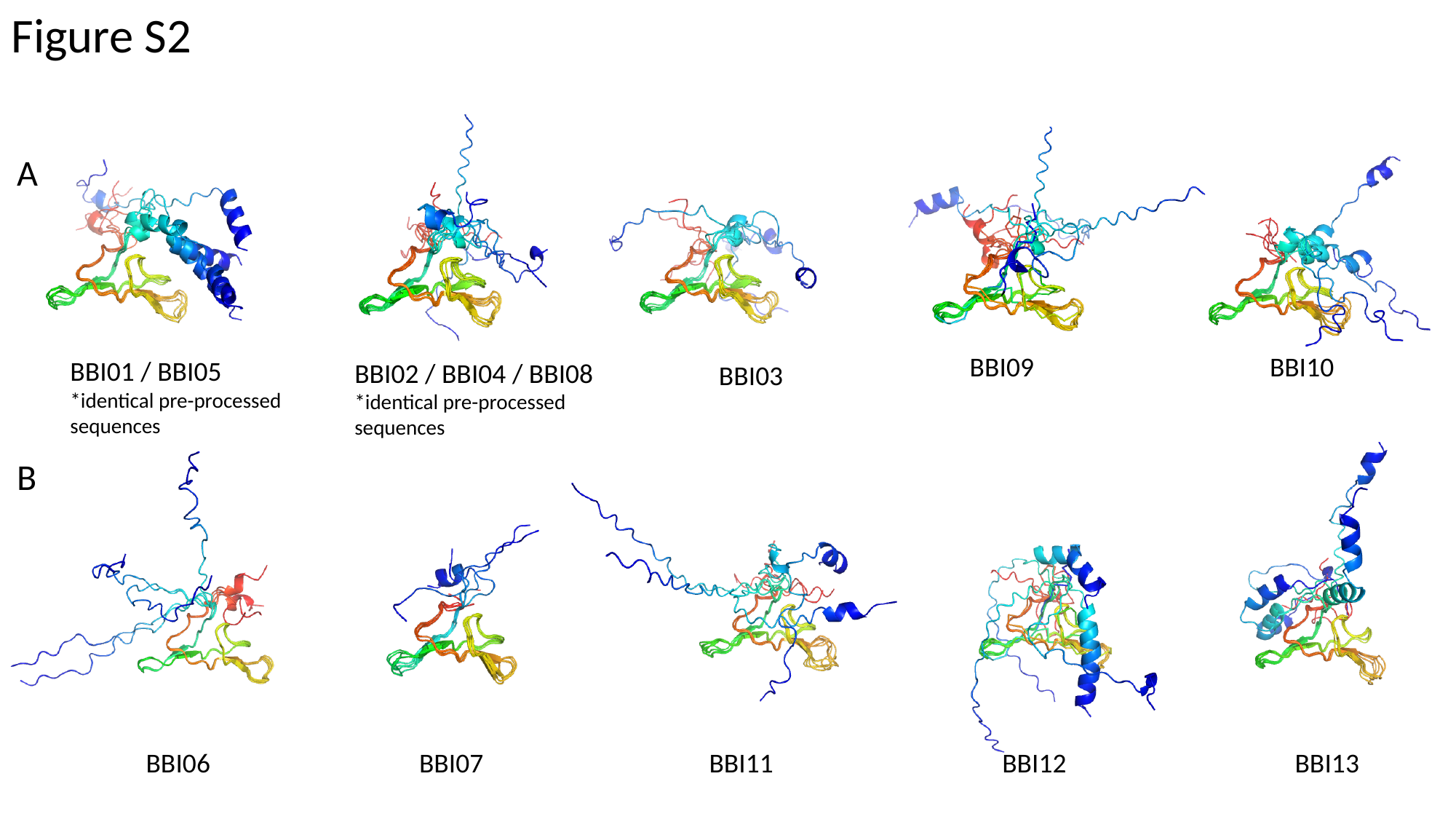

Figure S2
A
BBI09
BBI10
BBI01 / BBI05
*identical pre-processed sequences
BBI02 / BBI04 / BBI08
*identical pre-processed sequences
BBI03
B
BBI06
BBI07
BBI11
BBI12
BBI13

### Slide 3
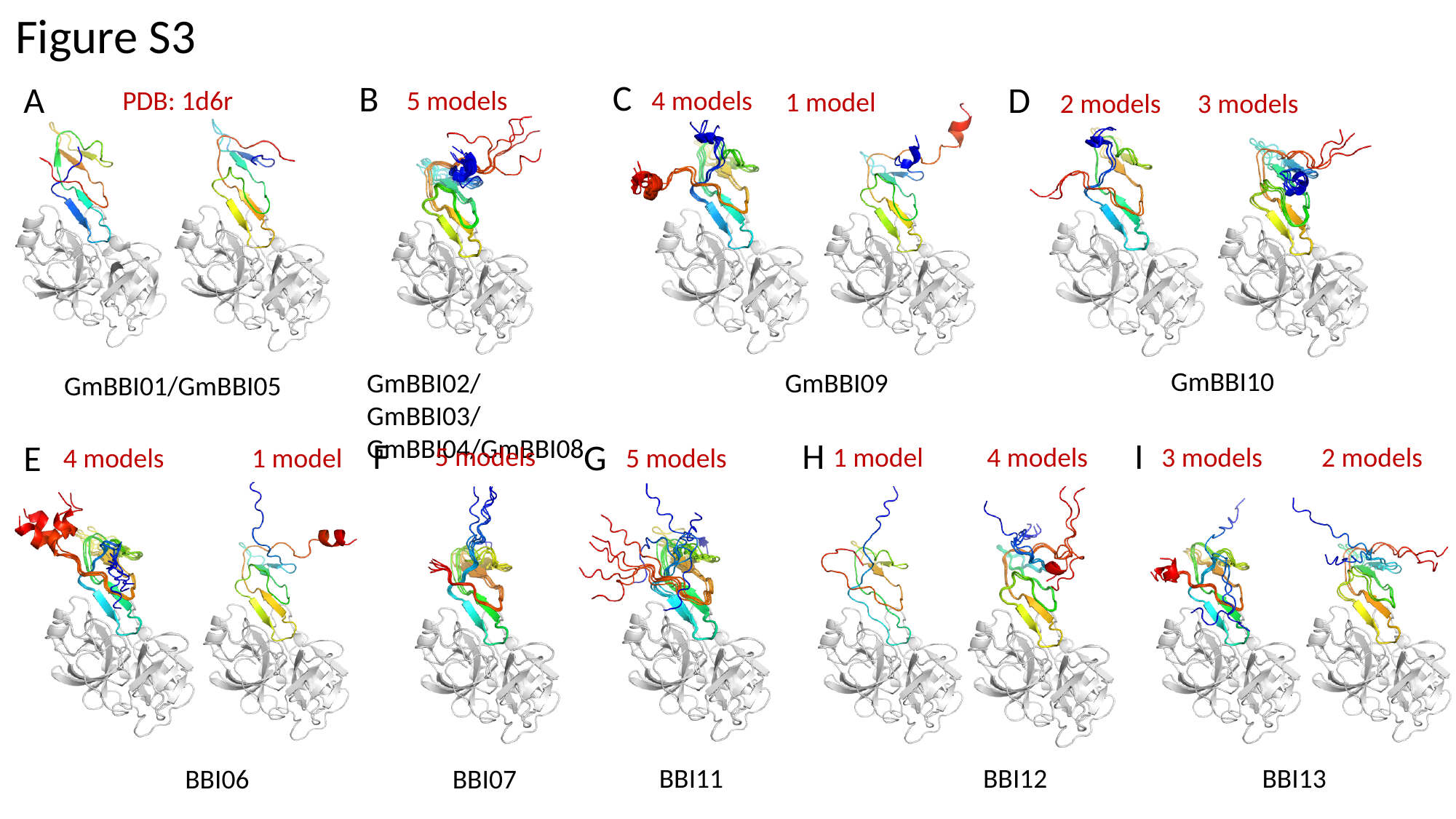

Figure S3
C
B
A
D
PDB: 1d6r
5 models
4 models
1 model
2 models
3 models
GmBBI10
GmBBI02/GmBBI03/GmBBI04/GmBBI08
GmBBI09
GmBBI01/GmBBI05
F
I
H
G
E
5 models
2 models
1 model
4 models
3 models
1 model
5 models
4 models
BBI11
BBI12
BBI13
BBI06
BBI07
BBI06
BBI12
BBI13

### Slide 4
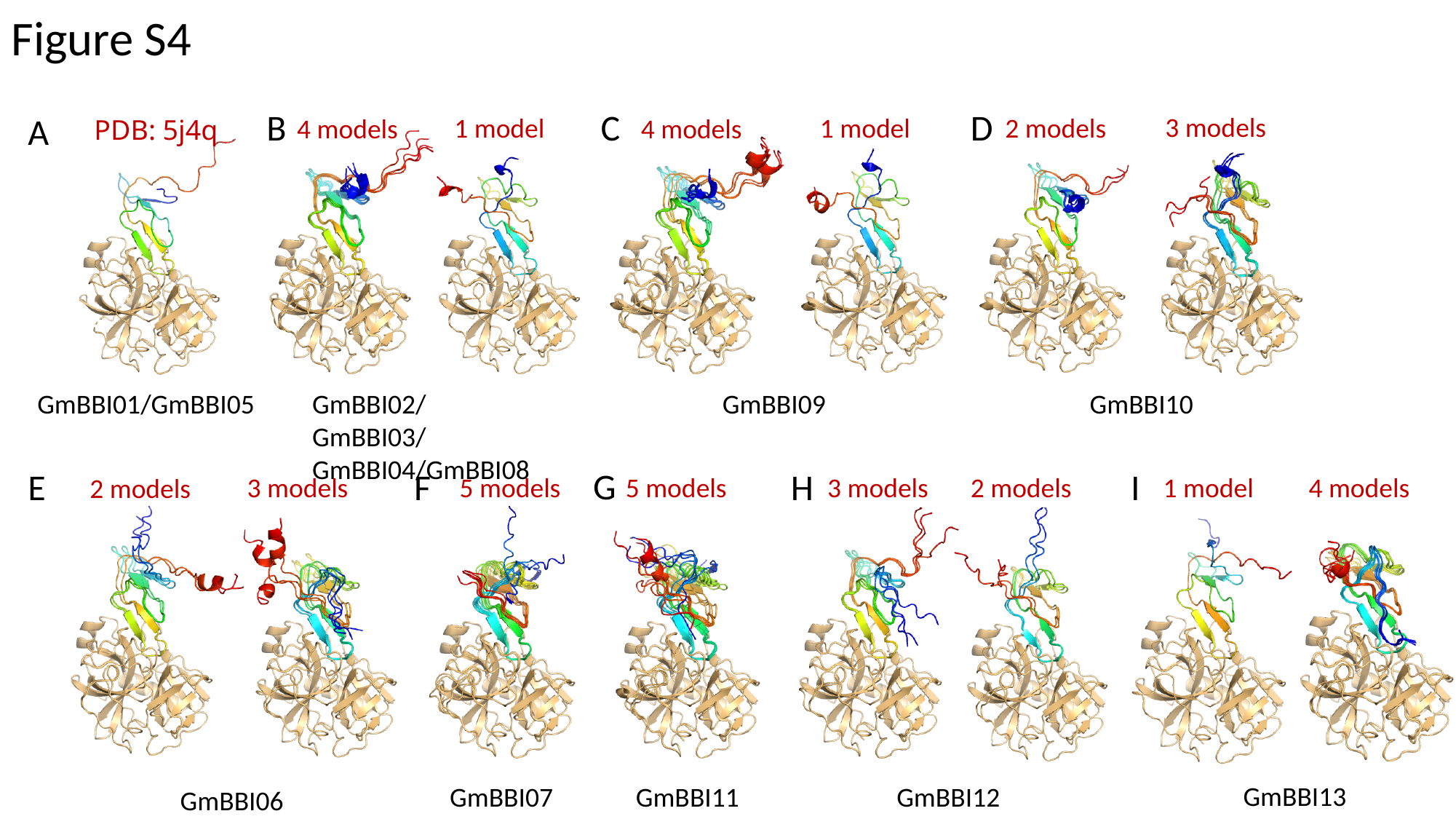

Figure S4
B
C
D
A
3 models
1 model
2 models
1 model
4 models
4 models
PDB: 5j4q
GmBBI01/GmBBI05
GmBBI09
GmBBI10
GmBBI02/GmBBI03/GmBBI04/GmBBI08
G
F
E
H
I
2 models
1 model
3 models
5 models
5 models
3 models
4 models
2 models
GmBBI13
GmBBI07
GmBBI11
GmBBI12
GmBBI06

### Slide 5
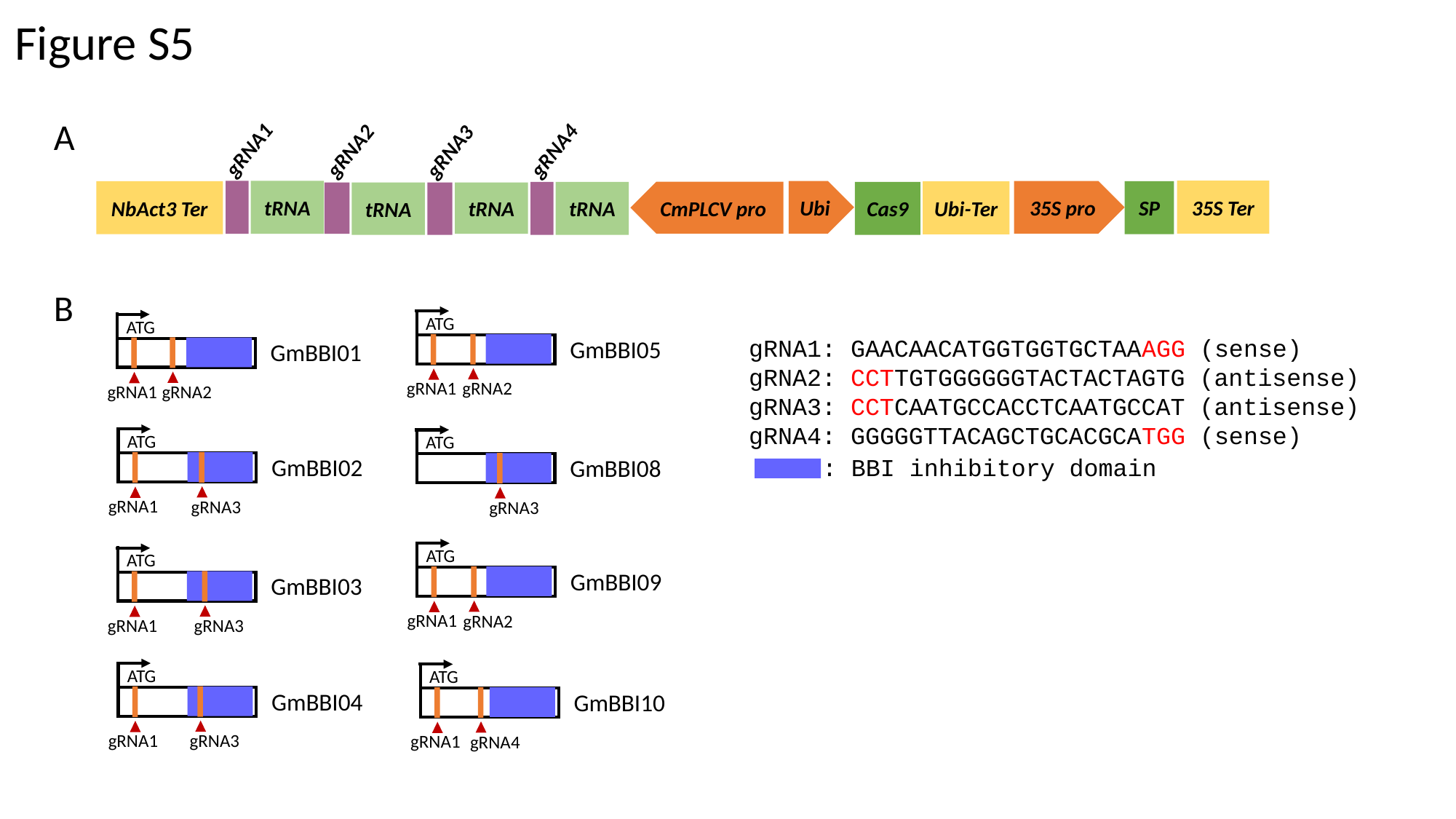

#### Figure S5
A
gRNA1
gRNA4
gRNA2
gRNA3
35S Ter
tRNA
Ubi
35S pro
SP
NbAct3 Ter
Ubi-Ter
tRNA
CmPLCV pro
Cas9
tRNA
tRNA
B
ATG
ATG
gRNA1: GAACAACATGGTGGTGCTAAAGG (sense)
gRNA2: CCTTGTGGGGGGTACTACTAGTG (antisense)
gRNA3: CCTCAATGCCACCTCAATGCCAT (antisense)
gRNA4: GGGGGTTACAGCTGCACGCATGG (sense)
GmBBI05
GmBBI01
gRNA1
gRNA2
gRNA1
gRNA2
ATG
ATG
GmBBI02
: BBI inhibitory domain
GmBBI08
gRNA1
gRNA3
gRNA3
ATG
ATG
GmBBI09
GmBBI03
gRNA1
gRNA2
gRNA1
gRNA3
ATG
ATG
GmBBI04
GmBBI10
gRNA1
gRNA3
gRNA1
gRNA4

### Slide 6
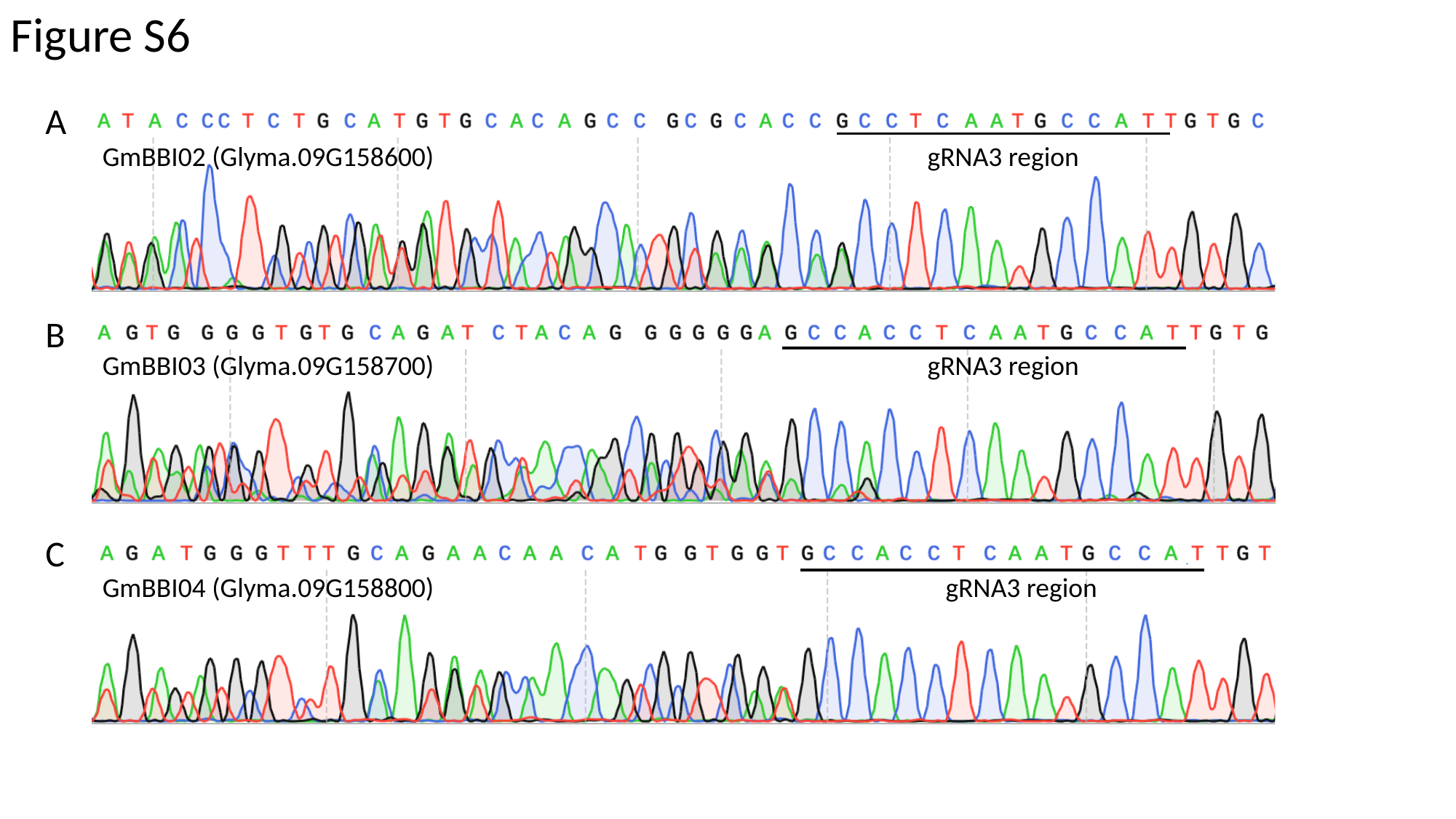

Figure S6
A
GmBBI02 (Glyma.09G158600)
gRNA3 region
B
GmBBI03 (Glyma.09G158700)
gRNA3 region
C
GmBBI04 (Glyma.09G158800)
gRNA3 region

### Slide 7
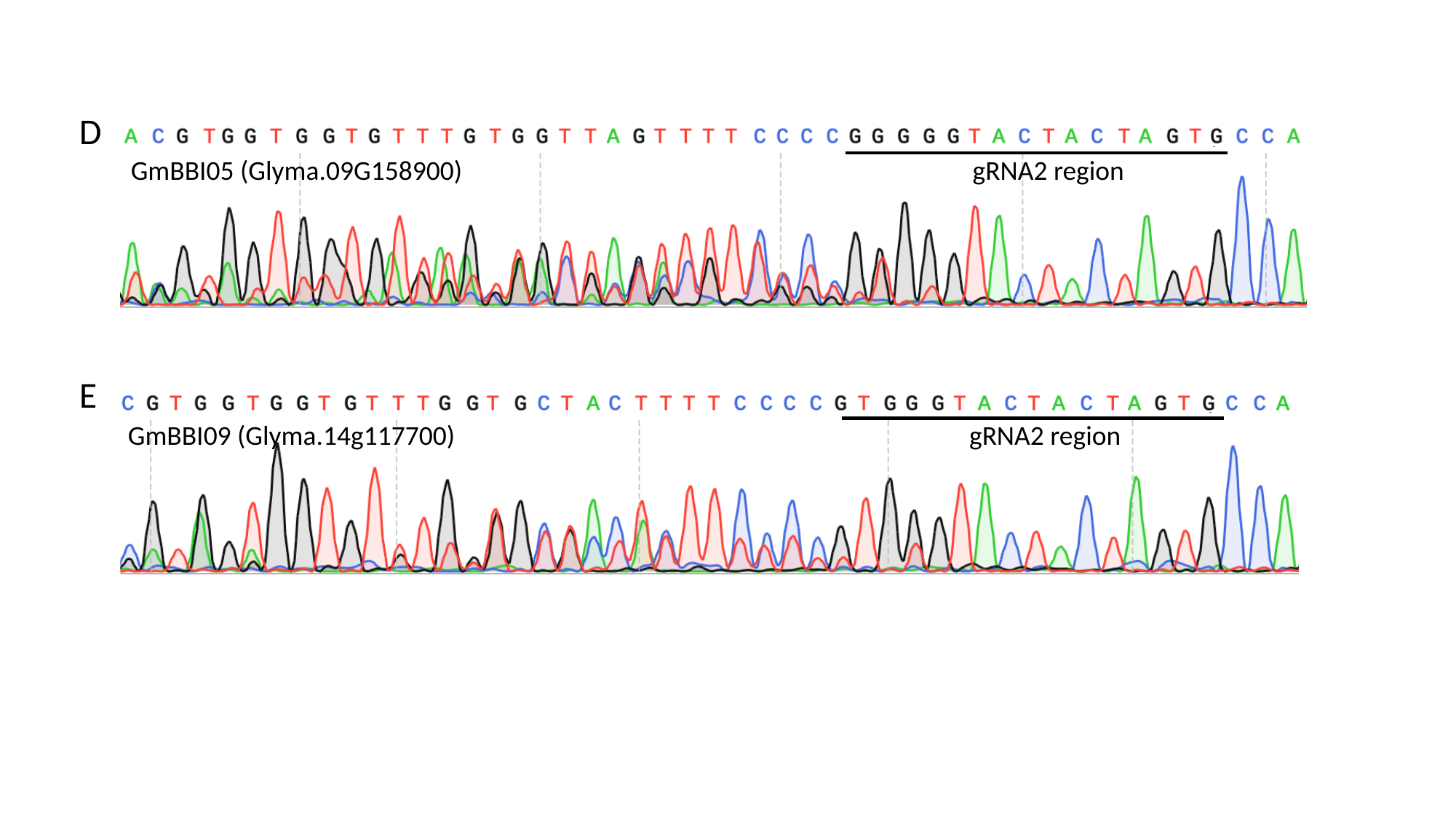

D
GmBBI05 (Glyma.09G158900)
gRNA2 region
E
GmBBI09 (Glyma.14g117700)
gRNA2 region

### Slide 8
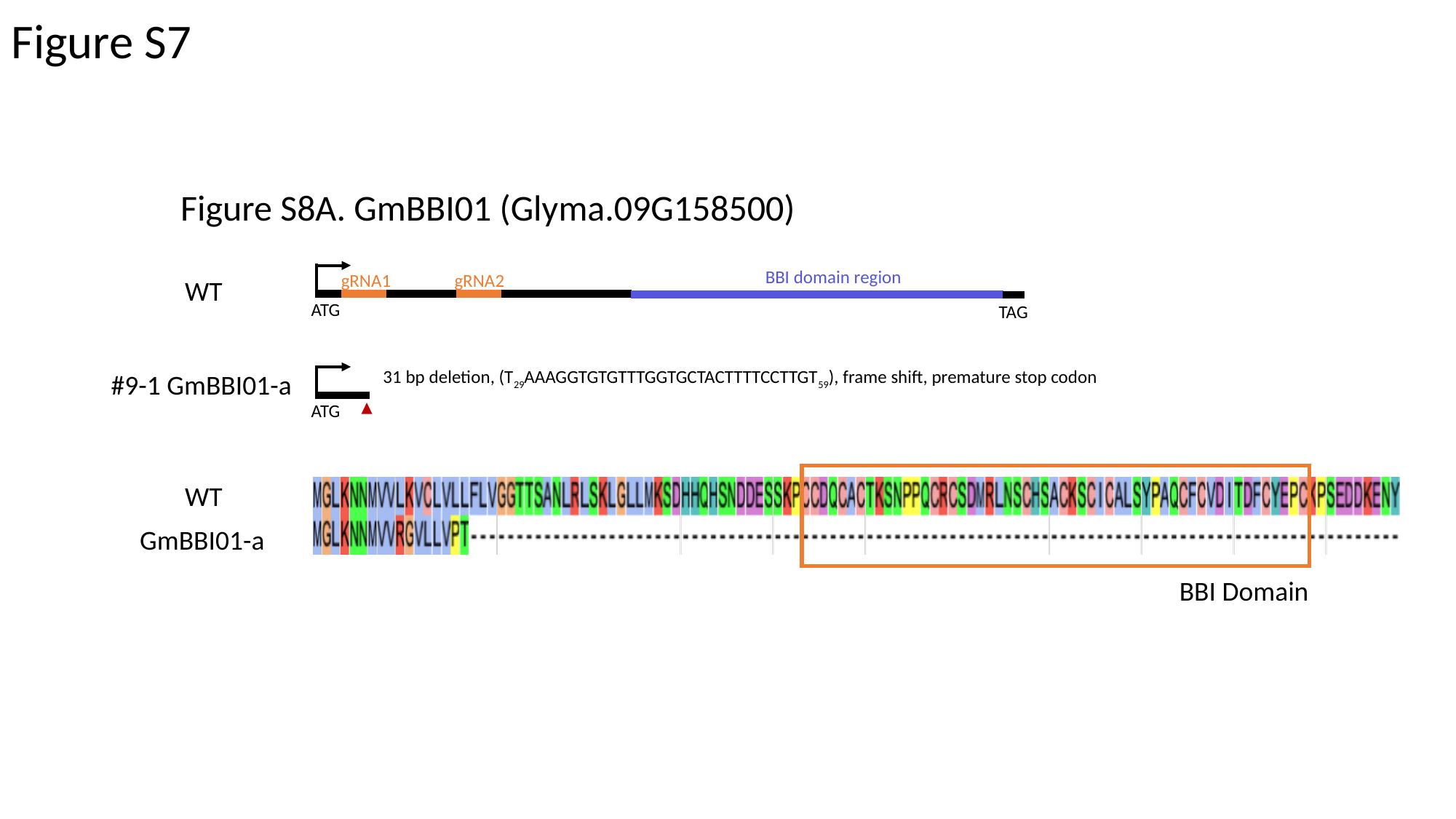

#### Figure S7
Figure S8A. GmBBI01 (Glyma.09G158500)
BBI domain region
gRNA1
gRNA2
WT
ATG
TAG
31 bp deletion, (T_29_AAAGGTGTGTTTGGTGCTACTTTTCCTTGT_59_), frame shift, premature stop codon
#9-1 GmBBI01-a
ATG
WT
GmBBI01-a
BBI Domain

### Slide 9
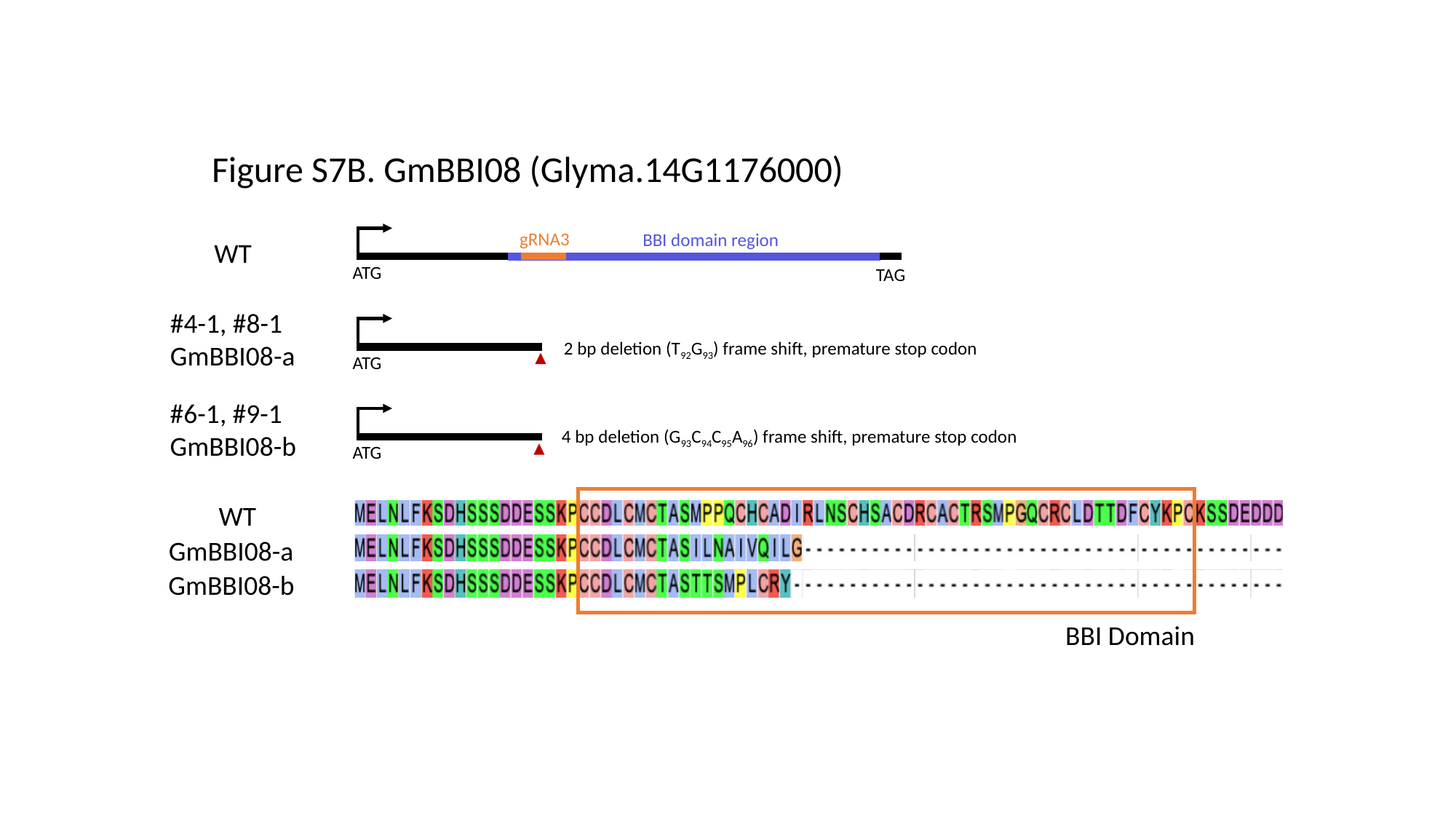

Figure S7B. GmBBI08 (Glyma.14G1176000)
gRNA3
BBI domain region
WT
ATG
TAG
#4-1, #8-1
GmBBI08-a
2 bp deletion (T_92_G_93_) frame shift, premature stop codon
ATG
#6-1, #9-1 GmBBI08-b
4 bp deletion (G_93_C_94_C_95_A_96_) frame shift, premature stop codon
ATG
WT
GmBBI08-a
GmBBI08-b
BBI Domain

### Slide 10
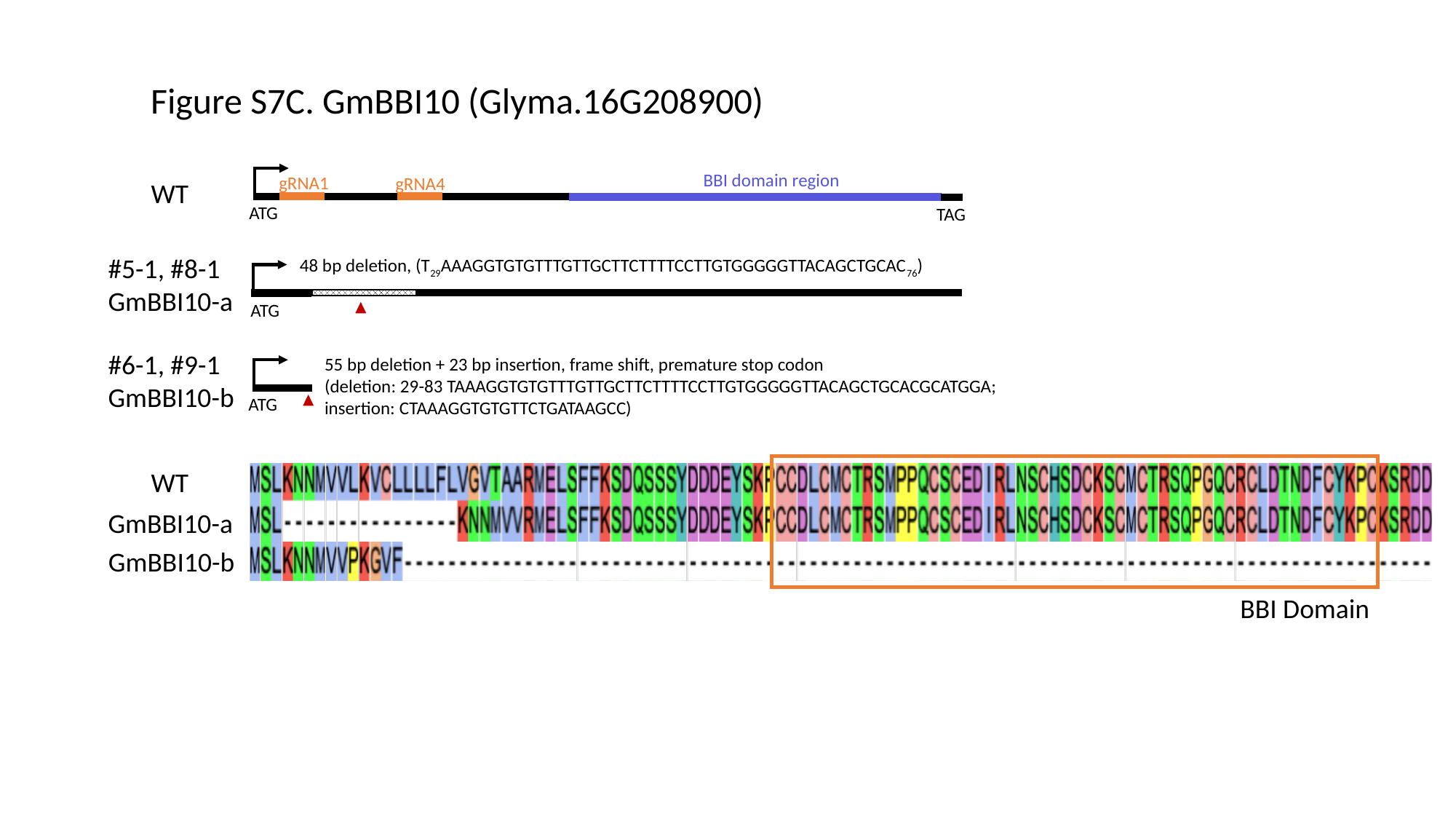

Figure S7C. GmBBI10 (Glyma.16G208900)
BBI domain region
gRNA1
gRNA4
WT
ATG
TAG
#5-1, #8-1 GmBBI10-a
48 bp deletion, (T_29_AAAGGTGTGTTTGTTGCTTCTTTTCCTTGTGGGGGTTACAGCTGCAC_76_)
ATG
#6-1, #9-1
GmBBI10-b
55 bp deletion + 23 bp insertion, frame shift, premature stop codon
(deletion: 29-83 TAAAGGTGTGTTTGTTGCTTCTTTTCCTTGTGGGGGTTACAGCTGCACGCATGGA; insertion: CTAAAGGTGTGTTCTGATAAGCC)
ATG
WT
GmBBI10-a
GmBBI10-b
BBI Domain
